# Distinct features in fish Bouncer proteins determine sperm-egg compatibility

**DOI:** 10.1101/2022.09.01.506233

**Authors:** Krista R.B. Gert, Karin Panser, Joachim Surm, Benjamin S. Steinmetz, Alexander Schleiffer, Yehu Moran, Fyodor Kondrashov, Andrea Pauli

## Abstract

All sexually reproducing organisms depend on fertilization to survive as species. Despite the importance of fertilization, the mechanisms that drive sperm-egg compatibility are poorly understood. In fish, the egg protein Bouncer is necessary for fertilization and is species-specific between medaka and zebrafish. Here, we investigate whether Bouncer is generally species-specific in fish and identify features mediating its medaka/zebrafish specificity. *In vitro* fertilization experiments using zebrafish and medaka show that Bouncer is not a general specificity factor. Instead, its homologs exhibit wide compatibility with sperm, in line with the pervasive purifying selection that dominates Bouncer’s evolution. We further uncover specific features of Bouncer— distinct amino acid residues and N-glycosylation patterns—that differentially influence the function of medaka and zebrafish Bouncer homologs and contribute to medaka/zebrafish specificity. This work reveals important themes central to understanding Bouncer’s function in sperm binding and clarifying the molecular requirements for Bouncer’s sperm interaction partner.

## Introduction

The reproductive success and continuation of all sexually reproducing species hinges upon fertilization, but our current understanding of gamete interaction and fusion is limited. Though studies over the past 20 years have identified several proteins essential for sperm-egg interaction, we lack a basic understanding of the molecular mechanisms and interaction partners for most factors. Indeed, only one mammalian sperm-egg interaction protein pair has been identified to date: IZUMO1 on sperm interacts with egg membrane-expressed JUNO to enable binding (*1*), (*2*). Though additional essential factors including Dcst1/2, Spaca6, TMEM95, FIMP, SOF1, and Bouncer have recently been discovered (*3*–*11*), their precise roles and interaction partners have yet to be described. Functional studies pinpointing important protein domains and molecular features of individual fertility factors are therefore crucial for understanding the mechanism of fertilization and can aid in the search for their interaction partners.

One important feature of many known gamete recognition proteins is species specificity (reviewed in (*12*)). Compatibility between gametes is critical for successful sperm-egg binding and fusion, but specificity is equally important for keeping fertilization restricted to members of a single species. Two classic examples of species-specific sperm-egg interactors are Bindin and EBR1 in sea urchin (*13*–*15*) and lysin and VERL in abalone (*16*–*19*). As broadcast spawners, these marine invertebrates rely on species-specific gamete interaction to avoid hybridization with other species that might be maladaptive. Because their eggs and sperm are at risk of encountering gametes from other abalone or sea urchin species within the same geographic range, a molecular block to cross-fertilization is therefore critical in the absence of other forms of pre-zygotic reproductive isolation.

In contrast, vertebrates such as fish and mammals have both anatomical and behavioral premating reproductive barriers that come into play prior to sperm-egg interaction. Despite premating reproductive isolation, mammals do have species-specific protein interactions between sperm and the zona pellucida (ZP), a glycoprotein matrix that surrounds the egg and is considered to act as a barrier to cross-species fertilization (*20, 21*). Studies exploiting the taxon specificity of human sperm binding to the ZP demonstrated that 32-34 amino acids at the N-terminus of one of the constituent ZP proteins, ZP2, is both necessary and sufficient for human sperm to bind to an otherwise mouse-derived ZP (*22, 23*). Though fish eggs do have a protective envelope surrounding the egg, the chorion, it contains a small opening, the micropyle, that allows direct contact of sperm with the egg membrane (*12, 24*). We previously showed that the egg membrane protein Bouncer (Bncr) is enriched at the micropyle and is not only required for sperm binding and entry in zebrafish eggs, but also is species-specific for medaka and zebrafish, two species that diverged ∼115-200 MYA, do not interbreed, and cannot cross-fertilize *in vitro* (*10, 25*). We found that expression of medaka Bncr in zebrafish *bncr*^−/-^ eggs enables fertilization by medaka sperm but not by zebrafish sperm (*10*). Similarly, expression of zebrafish Bncr in medaka eggs is sufficient for zebrafish sperm binding and fusion when these eggs are activated artificially after sperm addition (*26*). Importantly, Bncr provides specificity to the interaction of the egg and sperm membranes themselves, while the sperm-egg interactors described in marine invertebrates and mammals mediate specificity at the level of sperm interaction with the egg coat or ZP. Thus, in the absence of an outer layer conferring selectivity, Bncr may act analogously in allowing binding of only conspecific sperm to the egg membrane.

In this study, we investigated whether Bncr acts as a general species-specific factor in fish fertilization and sought to identify the molecular determinants in Bncr that mediate its specificity between zebrafish and medaka sperm.

## Results

### Medaka Bncra, but not Bncrb, is required on the medaka egg for fertilization

Bncr was originally identified and characterized in zebrafish (*10*), in which it exists as a single-exon gene (Fig. 1A, left). In medaka, however, it was unknown whether Bncr is also required for fertilization. Unlike the zebrafish locus, the medaka *bncr* locus gives rise to two Bncr splice isoforms whose mature domains are encoded by different exons, yet both adopt the characteristic Ly6/uPAR three-finger fold due to 8-10 invariant cysteines (*27*) (Fig. 1B). We therefore designated the two medaka proteins Bncra and Bncrb (Fig. 1A, right). The mature domain of medaka Bncra shares 38.8% identity with zebrafish Bncr and contains a predicted GPI anchor site and transmembrane domain on its C-terminus, like zebrafish Bncr. However, medaka Bncrb lacks these C-terminal features (Fig. 1A-B). Though absent in zebrafish, Bncrb is conserved in many other fish species (Fig. 1B; Suppl. Fig. 1A).

**Figure 1.**
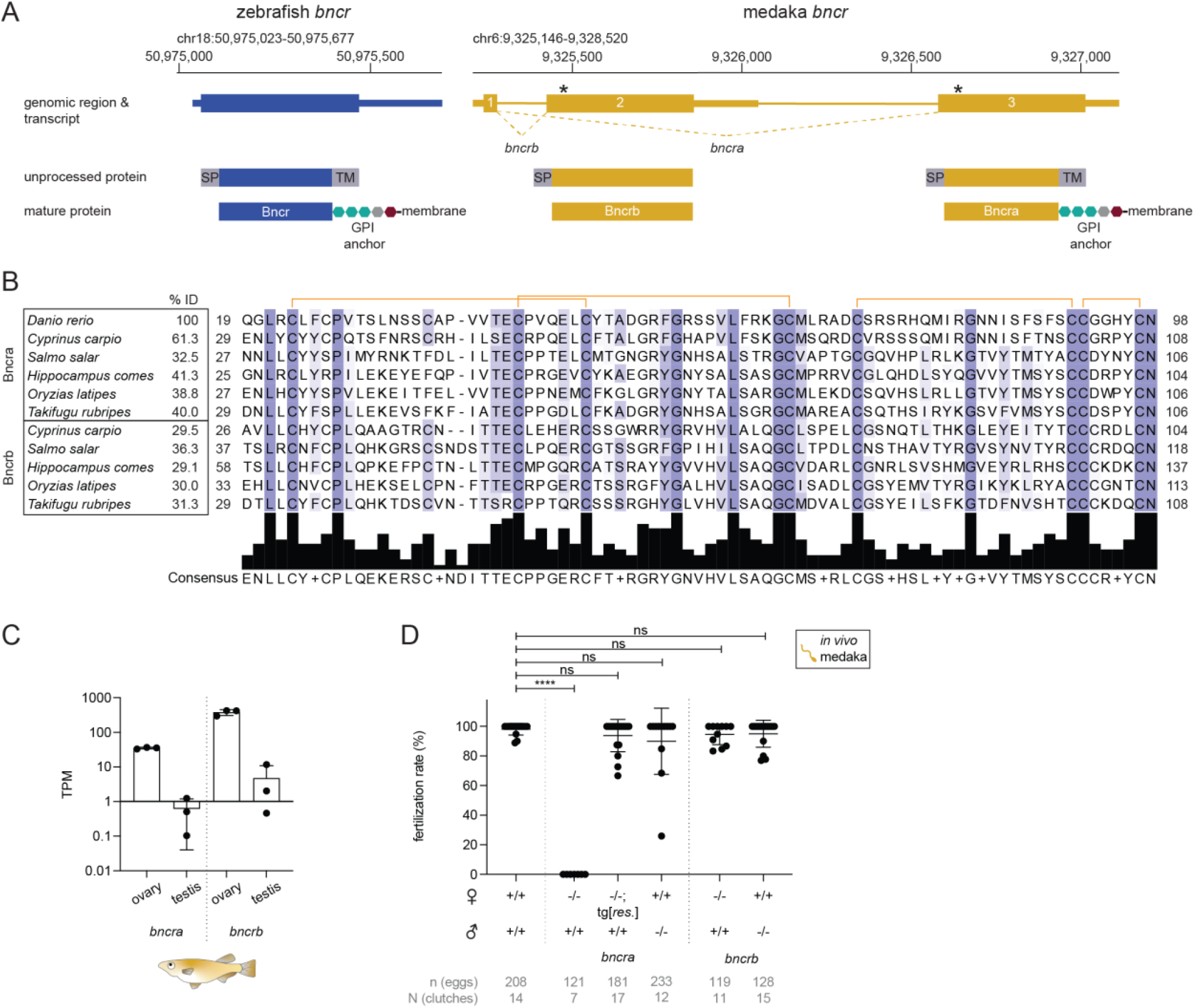
Medaka Bncra, but not Bncrb, is required on the medaka egg for fertilization. **(A)** Genomic regions and resulting transcripts and proteins of the *bncr* locus in zebrafish (blue; GRCz11/danRer11) and medaka (yellow; Ensembl 93: Jul 2018 (GRCh38.p12)). Zebrafish Bncr is encoded by a single-exon gene (NM_001365726.1) (left). The medaka *bncr* locus (ENSORLG00000004579) comprises three exons that are alternatively spliced to generate Bncra (exons 1 and 3; ENSORLT00000005754) and Bncrb (exons 1 and 2; ENSORLT00000005758). The location of the CRISPR-induced genomic deletions for medaka *bncra* and *bncrb* are indicated by asterisks. The gene structures are depicted with untranslated regions (thin rectangles) and coding sequences (thick rectangles). **(B)** Protein sequence alignment of the mature domains of Bncra and Bncrb from selected fish species (see Suppl. Data File 2). Note that seahorse Bncrb is only a predicted translation product from a genomic region. Purple shading indicates amino acids with at least 30% conservation. The percent amino acid sequence identity (% ID) within the mature domains is indicated. Disulfide bonds are indicated by orange brackets. **(C)** Expression values of *bncra* and *bncrb* transcripts in medaka ovary and testis based on RNA-seq (*28*). The Y-axis is plotted in log_10_ scale. TPM, transcripts per million. Means ± SD are indicated. **(D)** Quantification of *in vivo* fertilization rates from wild type and medaka *bncra* and *bncrb* mutants. Means ± SD are indicated. (Kruskal-Wallis test with Dunn’s multiple comparisons test: ****P < 0.0001 (wild-type ♀ x ♂ vs. *bncra*^−/-^ ♀ x wild-type ♂), P > 0.9999, ns (wild-type ♀ x ♂ vs. wild-type ♀ x *bncra*^−/-^ ♂, *bncra*^−/-^; tg[*bncra rescue*] ♀ x wild-type ♂, *bncrb*^−/-^ ♀ x wild-type ♂, and wild-type ♀ x *bncrb*^−/-^ ♂).

Because *bncra* and *bncrb* are both highly expressed in the medaka ovary (*28*) (Fig. 1C), we generated CRISPR/Cas9 mutants of both splice isoforms in medaka to investigate their potential roles in fertilization. Both the Bncra-specific mutation (5-nt deletion in exon 3) and the Bncrb-specific mutation (38-nt deletion in exon 2) (Fig. 1A) resulted in frameshifts leading to premature termination codons (Suppl. Data File 1). Crosses between *bncra*^−/-^ females and wild-type males revealed a similar phenotype as observed in zebrafish: Bncra-deficient eggs were neither activated nor fertilized, while *bncra*^−/-^ males were fertile when crossed to wild-type females (Fig. 1D). The sterility of *bncra*^−/-^ females could be rescued by an actin promoter-driven, GFP-tagged medaka *bncra* transgene (Fig. 1D). While zebrafish eggs undergo activation upon exposure to water, medaka eggs activate upon binding of sperm (*29, 30*).

Consistent with the defective sperm binding seen in *bncr*^−/-^ zebrafish eggs (*10*), medaka sperm fail to trigger activation of *bncra*^−/-^ eggs. In contrast to *bncra*^−/-^, *bncrb*^−/-^ females and males exhibited normal activation of eggs and fertility when crossed to wild-type fish (Fig. 1D). In line with its lack of predicted membrane anchorage, GFP-tagged Bncrb was secreted into the perivitelline space when expressed transgenically in zebrafish eggs, unlike membrane-localized, GFP-tagged Bncra (Suppl. Fig. 1B). Although Bncrb was not essential for fertility, it was possible that the presence of Bncra in *bncrb* mutants masked a potential function of Bncrb in fertilization. Because transgenic expression of medaka Bncra is sufficient to enable medaka sperm entry into a zebrafish egg (*10*), we assessed whether the same is true for Bncrb. To this end, we performed *in vitro* fertilization (IVF) experiments with zebrafish *bncr*^*-/-*^ eggs transgenically expressing GFP-tagged medaka Bncrb. However, neither zebrafish nor medaka sperm were able to fertilize these eggs (Suppl. Fig. 1C), demonstrating that Bncrb cannot rescue fertilization in zebrafish. Thus, medaka Bncra is required for fertilization in medaka and is homologous to zebrafish Bncr both in sequence and function, whereas Bncrb appears dispensable for this process.

### Medaka and zebrafish sperm are compatible with multiple Bncr homologs

Our observation that Bncra (hereafter Bncr) is species-specific between medaka and zebrafish (*10, 26*) prompted us to investigate whether other fish Bncr homologs also show evidence for species specificity by testing their compatibility with zebrafish and medaka sperm. For these experiments, we generated transgenic lines in *bncr*^−/-^ zebrafish expressing Bncr proteins from common carp (*Cyprinus carpio*), tiger tail seahorse (*Hippocampus comes*), and fugu (*Takifugu rubripes*), as well as a new *actin* promoter-driven medaka Bncr line. Because we previously observed a positive correlation between medaka *bncr* transcript level and fertilization rate (*10*), we used the *actin* promoter to drive expression of all transgenes in this study as it results in higher expression in the egg compared to the previously used *ubiquitin* promoter (*10, 31*).

As expected, the new *actin* promoter-driven medaka Bncr line had higher (more than an order of magnitude) transgene expression vs. the *ubiquitin* promoter-driven line (Suppl. Fig. 2A). In line with its increased expression, the *actin* promoter-driven medaka Bncr line had an average *in vitro* fertilization rate of 55.6% with medaka sperm (Fig. 2A) vs. only 5.7% for the *ubiquitin* promoter-driven line (*10*). Higher medaka Bncr expression in the egg also resulted in higher average *in vivo* and *in vitro* fertilization rates of 32.2% and 4.2%, respectively, with zebrafish sperm (Suppl. Fig 2B; Fig. 2A), indicating that the specificity for medaka sperm over zebrafish sperm can be partially overridden by medaka Bncr overexpression. Importantly, however, medaka sperm remain unable to fertilize zebrafish eggs overexpressing an *actin* promoter-driven zebrafish Bncr transgene expressed at similarly high levels as the *actin* promoter-driven medaka Bncr transgene (Fig. 2A; Suppl. Fig. 2A). This result suggests that the Bncr-mediated species specificity barrier is asymmetrical and could be governed by different features of Bncr for zebrafish vs. medaka sperm. Because all the tested transgenes were expressed on zebrafish eggs, there might also be intrinsic bias for zebrafish sperm given conspecificity of the egg in general. However, other work has shown that zebrafish sperm are better able to fertilize wild-type medaka eggs (2% on average) than medaka sperm can fertilize wild-type zebrafish eggs *in vitro* (0%) (*26*), further supporting asymmetry in specificity between these sperm-Bncr interactions.

**Figure 2.**
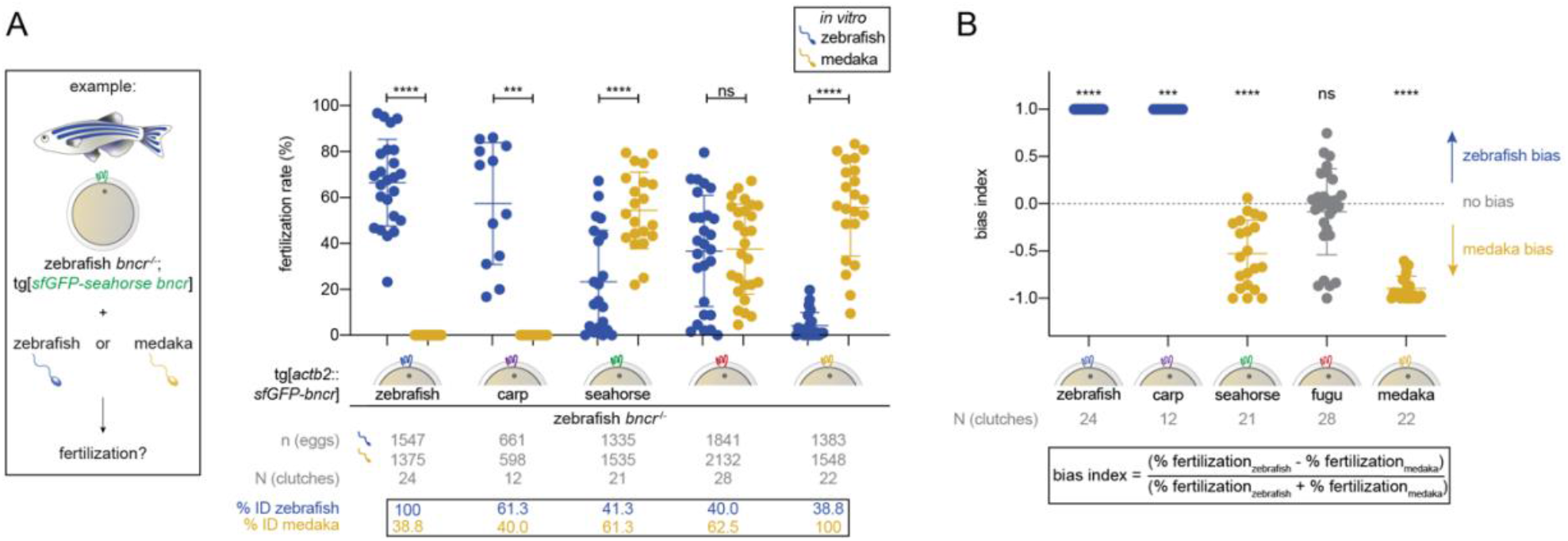
Medaka and zebrafish sperm are compatible with multiple Bncr homologs. **(A)** Experimental setup for performing comparative medaka/zebrafish IVF with transgenic zebrafish *bncr*^*-/-*^ eggs expressing different fish Bncr homologs (left). IVF data obtained from transgenic zebrafish *bncr*^*-/-*^ lines expressing either zebrafish, carp, seahorse, fugu, or medaka Bncr with medaka vs. zebrafish sperm (right). Means ± SD are indicated. (Wilcoxon matched-pairs signed rank test with method of Pratt: ****P < 0.0001 (zebrafish vs. medaka sperm with zebrafish, seahorse, and medaka Bncr), ***P = 0.0005 (carp), P = 0.6277, ns (fugu)). **(B)** Plot of the bias index values derived from the IVF data in Fig. 2A. The formula for the bias index is shown. (Wilcoxon signed rank test vs. theoretical median of 0 with method of Pratt: ****P < 0.0001 (zebrafish, seahorse, and medaka), ***P = 0.0005 (carp), P = 0.5824, ns (fugu)).

Contrary to the hypothesis that Bncr is a general species specificity factor in fish, zebrafish sperm were compatible with carp, seahorse, and fugu Bncr *in vivo* and *in vitro* (Suppl. Fig. 2B; Fig. 2A). Moreover, while medaka sperm failed to fertilize carp Bncr-expressing zebrafish eggs, they fertilized seahorse and fugu Bncr-expressing zebrafish eggs with average fertilization rates of 54.4% and 36.7% *in vitro* (Fig. 2A). Thus, seahorse and fugu Bncr are compatible with both zebrafish and medaka sperm. To assess the relative bias of the tested Bncr proteins for zebrafish vs. medaka sperm, we calculated the bias index (Fig. 2B) using the IVF data (Fig. 2A) for each line. While zebrafish and carp Bncr strictly favor zebrafish sperm, seahorse and medaka Bncr display bias for medaka sperm (Fig. 2B). Interestingly, fugu Bncr does not exhibit bias for either sperm (Fig. 2B). This result suggests that the features required for successful interaction with both medaka and zebrafish sperm can coexist in the same Bncr protein and are therefore not mutually exclusive.

### Medaka/zebrafish Bncr chimeras reveal specificity determinants in fingers 2 and 3

To investigate which parts of the Bncr protein (referred to as “fingers” given Bncr’s three-finger fold (*10, 27*)) confer medaka/zebrafish specificity, we generated a set of transgenic zebrafish lines that express medaka/zebrafish Bncr chimeras in the zebrafish *bncr*^−/-^ background. These chimeras comprise eight different combinations of fingers as well as the upper (top, all three fingers excluding the base) and lower (base) regions of medaka and zebrafish Bncr (Fig. 3A; Suppl. Data File 3), such that the invariant cysteines were the boundary between the “top” and “base”.

**Figure 3.**
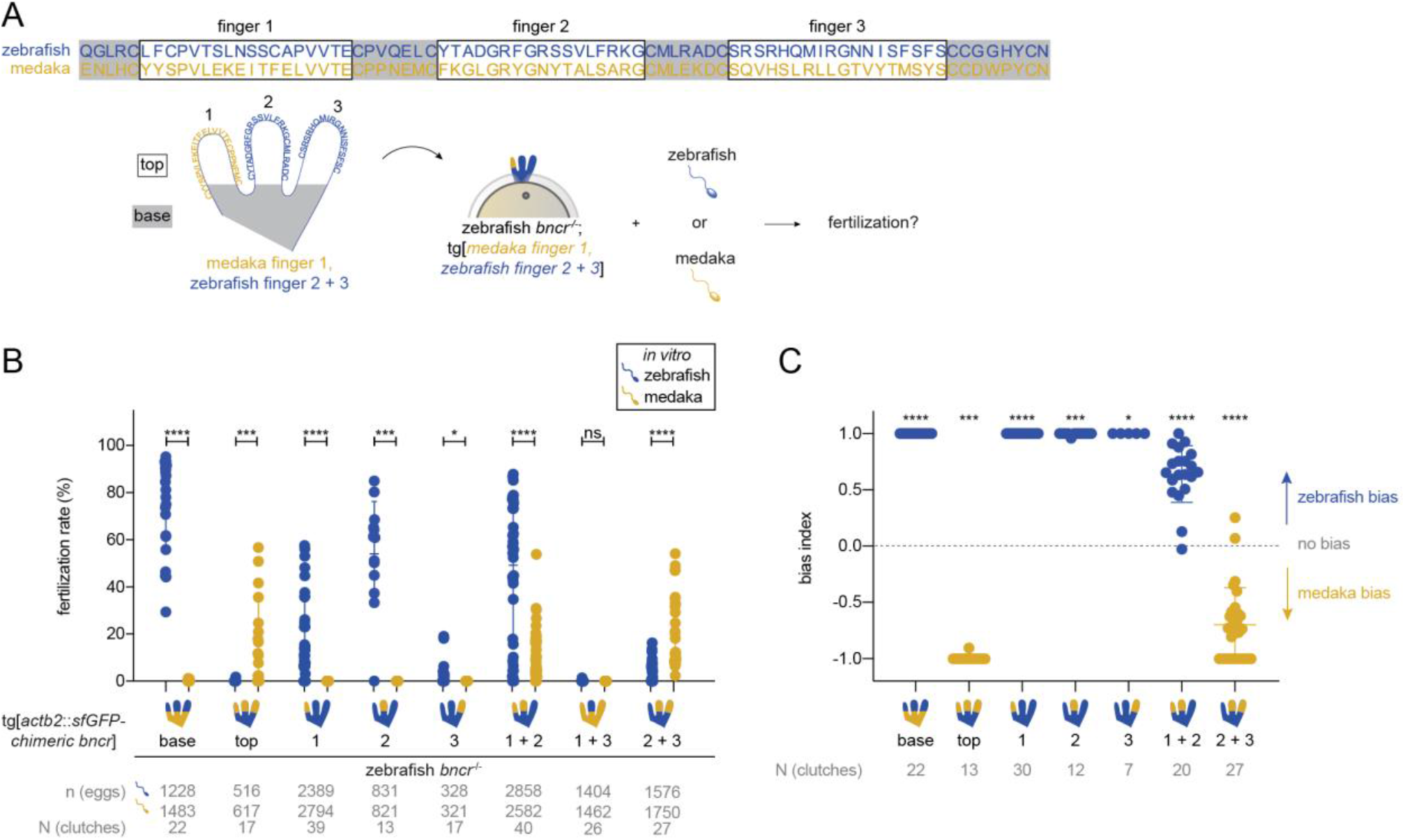
Medaka/zebrafish Bncr chimeras reveal specificity determinants in fingers 2 and 3. **(A)** Zebrafish (blue) and medaka (yellow) mature Bncr protein sequence alignment and schematic of the Bncr protein fold. Fingers are labeled 1, 2, and 3 and correspond to the amino acids in boxes in the protein sequence alignment. Note that each finger is bounded by cysteine residues that keep disulfide bridges intact. “Top” and “base” are indicated. **(B)** Comparative medaka/zebrafish IVF data with Bncr chimera lines. Means ± SD are indicated. (Wilcoxon matched-pairs signed rank test with method of Pratt: ****P < 0.0001 (zebrafish vs. medaka sperm with chimeras 1, 1+2, 2+3, and base), ***P = 0.0005 (2), ***P = 0.0002 (top), *P = 0.0156 (3); P > 0.9999, ns (1+3)). **(C)** Plot of the bias index values derived from the IVF data in Fig. 3B. Bias could not be calculated for data pairs for which the fertilization rate with both sperm was equal to 0. Means ± SD are indicated. (Wilcoxon signed rank test vs. theoretical median of 0 with method of Pratt: ****P < 0.0001 (1, 1+2, 2+3, base), ***P = 0.0005 (2), ***P = 0.0002 (top), *P = 0.0156 (3)).

With this approach, we systematically tested the role of each finger or combination of fingers for compatibility with medaka vs. zebrafish sperm in IVF experiments (Fig. 3A). Changing only the “top” but not the “base” to the medaka sequence enabled fertilization by medaka sperm and abrogated fertilization by zebrafish sperm, revealing that the species specificity determinants are encoded within the upper regions of the three fingers (Fig. 3B). Single medaka finger substitutions were not sufficient to rescue fertilization with medaka sperm. Changing finger 3 to medaka greatly decreased fertilization rates with zebrafish sperm *in vitro*, suggesting a role for finger 3 in mediating specificity (Fig. 3B), though fertilization rates *in vivo* remained high (72.7% on average) (Suppl. Fig. 3A). Combinations of medaka fingers 1 + 2 and 2 + 3 were compatible with both species’ sperm. Combining medaka fingers 1 + 3 failed to rescue fertilization with either sperm *in vitro* (Fig. 3B) despite low *in vivo* fertilization rates (2.5% on average) with zebrafish sperm (Suppl. Fig. 3A) and expression on the egg membrane (Suppl. Fig. 3B). This chimera’s inability to rescue fertilization with either species’ sperm precluded it from bias calculation. Though compatible with both species’ sperm, the medaka finger 1 + 2 chimera showed a clear bias for zebrafish sperm (Fig. 3C). Bias for medaka over zebrafish sperm was evident only upon changing fingers 2 + 3 together or all three (top) to the medaka sequence (Fig. 3C). This data demonstrates a requirement for medaka finger 2 in addition to either finger 1 or 3 for medaka sperm compatibility, with finger 3 having a stronger effect in shifting bias toward medaka sperm. In addition, chimeras containing zebrafish finger 3 maintain a bias for zebrafish sperm, further underscoring a role for finger 3 in determining species specificity. These results hint toward clarifying the asymmetric requirements in Bncr for medaka vs. zebrafish sperm: while medaka require features in both fingers 2 + 3 for specificity, only finger 3 is required for zebrafish specificity.

### Ancestral Bouncer states reveal a positively selected, Oryzias-specific change that hampers zebrafish sperm compatibility

To identify more precisely the features within medaka and zebrafish Bncr that underlie the incompatibility between these two species’ gametes, we took an evolutionary approach. First, using fish Bncr phylogeny, we predicted ancestral states of the Bncr protein (see Methods) that contain the predicted changes undergone between the zebrafish and medaka homologs (Fig. 4A-B; Suppl. Data File 3). To identify when in Bncr’s evolutionary history incompatibility with zebrafish sperm may have arisen and which amino acid changes caused this, we generated transgenic lines in the zebrafish *bncr*^*-/-*^ background expressing the predicted ancestral states of Bncr between seahorse and fugu Bncr and tested them for fertility with zebrafish and medaka sperm. In line with the dual compatibility observed for seahorse and fugu Bncr, ancestral states at nodes A-D (the same sequence was predicted for these four nodes), E, and G exhibited compatibility for both species’ sperm (Fig. 4B-C). Nodes A-D and G, however, rescued poorly with both zebrafish and medaka sperm despite expression at the egg membrane and the ability to rescue fertilization *in vivo* with zebrafish sperm (Suppl. Fig. 4A-B), suggesting that these Bncr states contain features detrimental for interaction with both sperm. These nodes were excluded from bias calculation due to their overall inefficient rescue. While the ancestral Bncr at node E showed similar compatibility with both zebrafish and medaka sperm *in vitro*, a clear bias for medaka sperm was observed at node F which immediately precedes the *Oryzias* (medaka) genus clade (Fig. 4C-D), pointing toward the presence of an *Oryzias*-specific change that hinders zebrafish compatibility.

**Figure 4.**
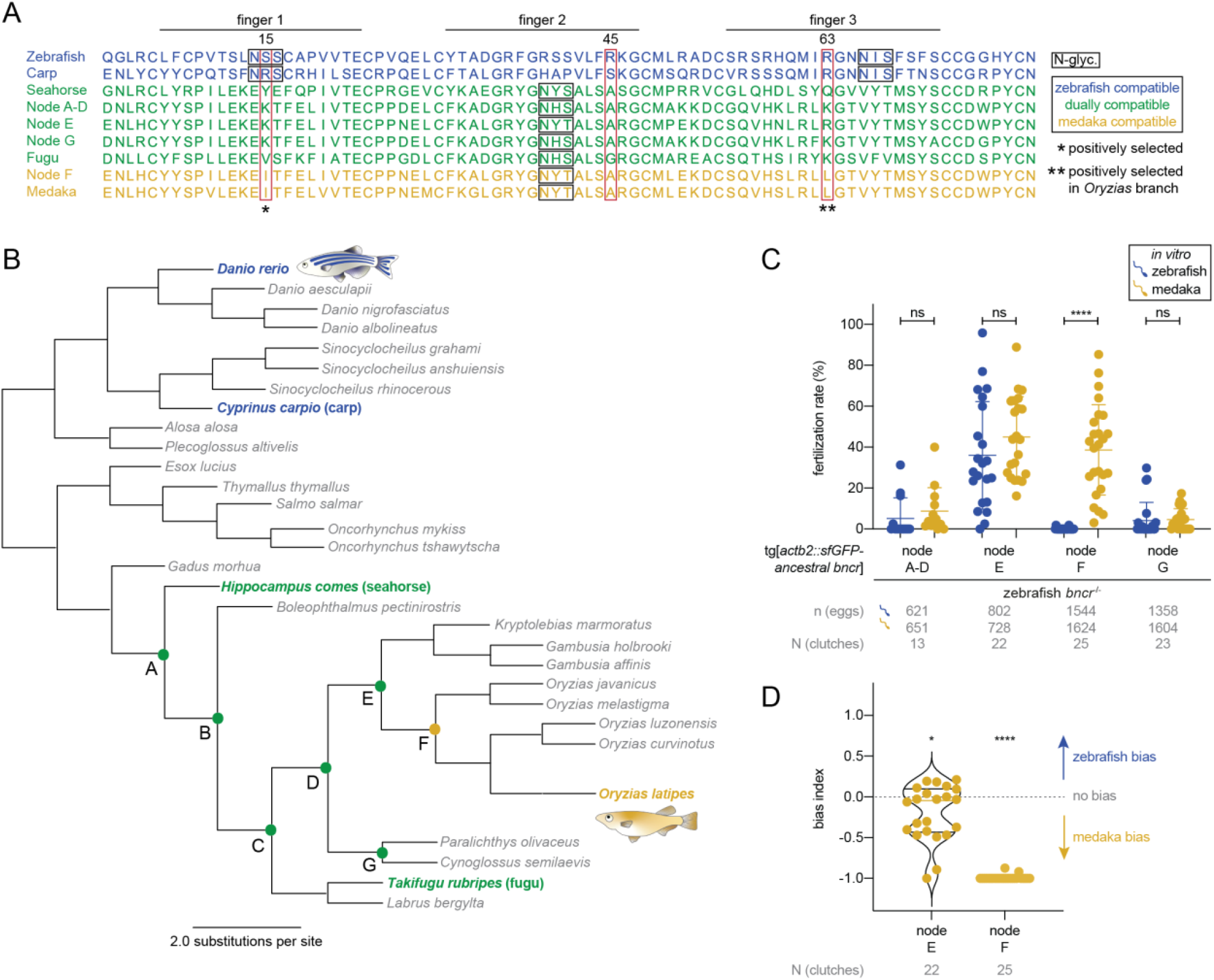
Ancestral Bncr states reveal a positively selected, *Oryzias*-specific change that hampers zebrafish sperm compatibility. **(A)** Protein sequence alignment of fish Bncr homologs and predicted ancestral states of Bncr. Fingers 1, 2, and 3 are indicated. Zebrafish-compatible sequences are blue, medaka-compatible sequences are yellow, and dually compatible sequences are green. Amino acid numbering is based on mature Bncr sequences. Red rectangles demarcate sites 15, 45, and 63 which were tested individually and in combination for their role in species specificity, while N-glycosylation sites are marked with a black rectangle (see Fig. 5F). The two positively selected sites are highlighted with asterisks. **(B)** Phylogenetic tree of the predicted Bncr ancestral states according to fish phylogeny. Tested nodes (A-G) are marked with a closed circle and colored according to compatibility as in Figure 4A. Nodes A-D were predicted to have the same sequence and are therefore equivalent. **(C)** Comparative medaka/zebrafish IVF data from the tested Bncr ancestral states. Means ± SD are indicated. (Wilcoxon matched-pairs signed rank test with method of Pratt: ****P < 0.0001 (zebrafish vs. medaka sperm with node F); P = 0.0830, ns (node A-D); P = 0.0917, ns (node E); P = 0.3591, ns (node G)). **(D)** Plot of bias index values derived from the IVF data pairs from node E and node F in Fig. 4C. Means ± SD are indicated. (Wilcoxon signed rank test vs. theoretical median of 0 with method of Pratt: ****P < 0.0001 (node F), *P = 0.0391 (node E)).

In a second evolutionary approach, we performed positive selection analysis of the mature domain of fish Bncr proteins (see Methods). Although Bncr’s evolution is dominated by pervasive purifying (negative) selection, indicating pressure to conserve the amino acid sequence and thereby preserve binding interactions, the *Oryzias* Bncr branch specifically was found to be under episodic diversifying (positive) selection, indicating the presence of medaka-specific changes that might underlie the higher Bncr specificity observed for these fish (Fig. 4A; Suppl. Fig. 4C; Suppl. Data File 4). Overall, two positively selected sites were detected. Site 15 (Ser in zebrafish; Ile in medaka) in finger 1 had signatures of pervasive diversifying selection throughout the phylogeny of tested fish Bncr proteins, while site 63 (Arg in zebrafish; Leu in medaka) in finger 3 had evidence of episodic diversifying selection specifically in the *Oryzias* lineage (Fig. 4A; Suppl. Fig. 4C; Suppl. Data File 4). Both positively selected sites differed between the ancestral Bncr sequences at nodes E and F, concomitant with a switch in bias from zebrafish to medaka sperm (Fig. 4A, D), suggesting a possible contribution to the observed species specificity.

Based on our evolutionary analyses, we tested the contribution of sites 15 and 63 in determining medaka/zebrafish specificity. Moreover, given the chimera data that implicated finger 2 in medaka sperm compatibility, we further compared medaka-compatible vs. incompatible Bncr sequences and identified site 45 as another candidate that might contribute to specificity (Arg in zebrafish; Ala or Gly in all medaka-compatible sequences) (Fig. 4A; Suppl. Data File 3). Based on the AlphaFold structural predictions (*32*) of zebrafish and medaka Bncr, the two arginines in sites 45 and 63 may together form a positively charged patch in zebrafish Bncr that is absent in medaka Bncr (Fig. 5A-B). We hypothesized that this positively charged patch may either be unfavorable for medaka sperm or beneficial for zebrafish sperm interaction.

**Figure 5.**
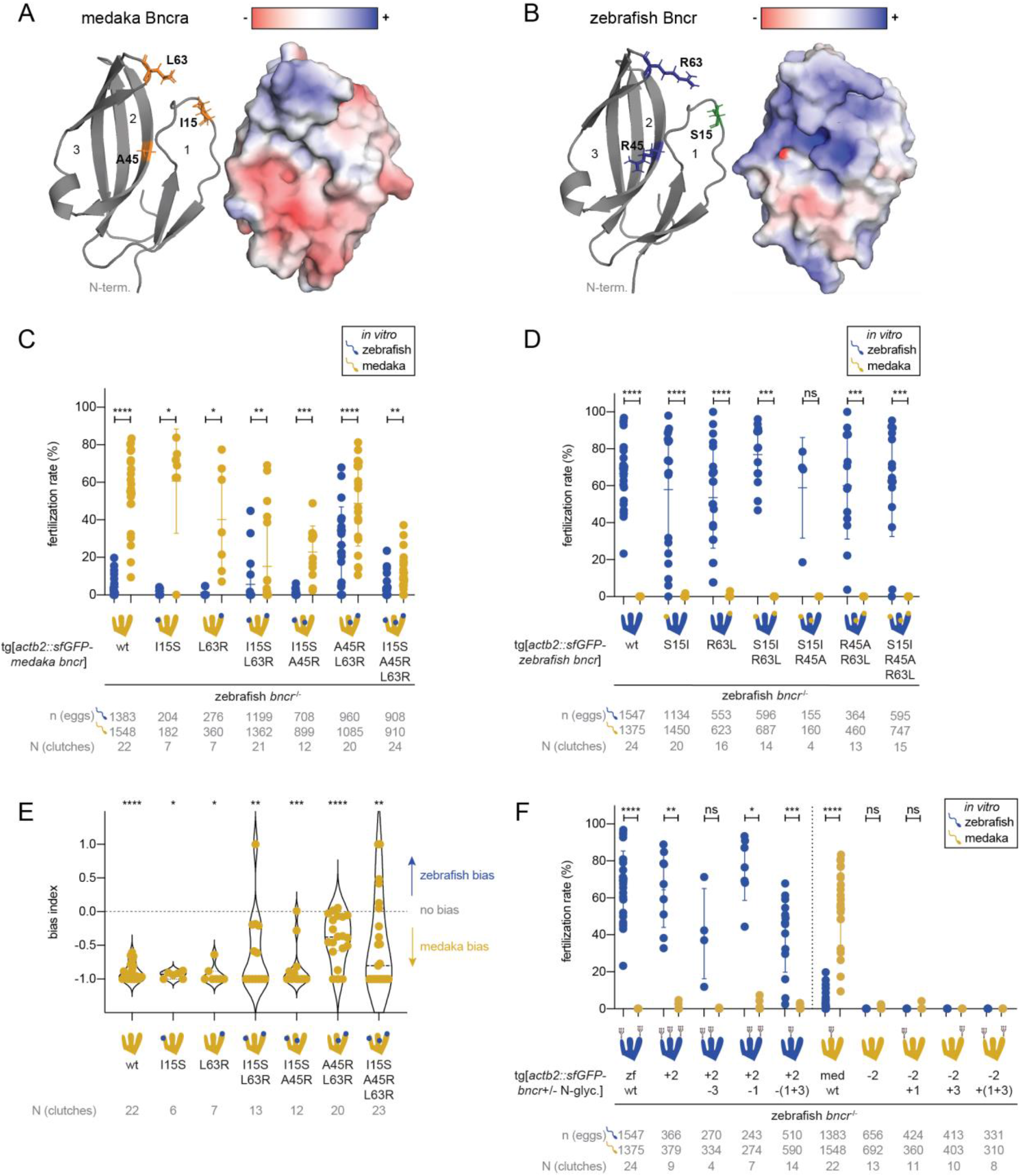
Zebrafish sperm favor a positively charged Bncr surface, while medaka sperm require finger 2 N-glycosylation for compatibility. **(A)** AlphaFold-predicted model of medaka Bncr (cartoon, left; surface representation depicting electrostatics, right). Amino acids that were mutated are indicated in the model as sticks and are colored orange to indicate hydrophobic side chains. **(B)** AlphaFold-predicted model of zebrafish Bncr (cartoon, left; surface representation depicting electrostatics, right). Amino acids that were mutated are indicated in the model as sticks; positively charged side chains are blue, while the polar side chain is green. **(C)** Medaka/zebrafish IVF with medaka Bncr constructs, in which individual amino acids or combinations thereof were substituted for the corresponding amino acid(s) in zebrafish Bncr. Means ± SD are indicated. (Wilcoxon matched-pairs signed rank test with method of Pratt: ****P < 0.0001 (zebrafish vs. medaka sperm with medaka Bncr and medaka Bncr A45R, L63R), ***P = 0.0010 (medaka Bncr I15S, A45R), **P = 0.0039 (medaka Bncr I15S, L63R), **P = 0.0029 (medaka Bncr I15S, A45R, L63R), *P = 0.0156 (medaka Bncr L63R), *P = 0.0312 (medaka Bncr I15S)). **(D)** Medaka/zebrafish IVF with zebrafish Bncr constructs, in which individual amino acids or combinations thereof were substituted for the corresponding amino acid(s) in medaka Bncr. Means ± SD are indicated. (Wilcoxon matched-pairs signed rank test with method of Pratt: ****P < 0.0001 (zebrafish vs. medaka sperm with zebrafish Bncr, zebrafish R63L, and zebrafish S15I), ***P = 0.0001 (zebrafish S15I, R63L and zebrafish S15I, R45A, R63L), ***P = 0.0002 (zebrafish R45A, R63L), P = 0.1250, ns (zebrafish S15I, R45A)). **(E)** Plot of bias index values derived from the IVF data in Fig. 5C. Bias could not be calculated for data pairs for which the fertilization rate with both sperm was equal to 0. Means ± SD are indicated. (Wilcoxon signed rank test vs. theoretical median of 0 with method of Pratt: ****P < 0.0001 (zebrafish vs. medaka sperm with medaka Bncr and medaka Bncr A45R, L63R), ***P = 0.0010 (medaka Bncr I15S, A45R), **P = 0.0073 (medaka Bncr I15S, L63R), **P = 0.0042 (medaka Bncr I15S, A45R, L63R), *P = 0.0312 (medaka Bncr I15S), *P = 0.0156 (medaka Bncr L63R)). **(F)** Medaka/zebrafish IVF experiments to assess the importance of N-glycosylation in Bncr’s species-specificity. IVF with zebrafish Bncr N-glycosylation site variants (left); IVF with medaka Bncr N-glycosylation site variants (right). Means ± SD are indicated. (Wilcoxon matched-pairs signed rank test with method of Pratt: ****P < 0.0001 (medaka vs. zebrafish sperm with medaka Bncr and zebrafish Bncr), ***P = 0.0001 (zebrafish Bncr -(glyc 1 + 3) + glyc 2), **P = 0.0039 (zebrafish Bncr + glyc 2), *P = 0.0156 (zebrafish Bncr + glyc 2, - glyc 1), P = 0.2500, ns (medaka Bncr - glyc 2), P = 0.1250, ns (zebrafish Bncr + glyc 2, - glyc 3), and P < 0.9999, ns (medaka Bncr - glyc 2, + glyc 1)). For the constructs medaka Bncr - glyc 2, + glyc 3 and medaka Bncr - glyc 2, +(glyc 1 + 3), p values could not be calculated as all data points are 0.

Using transgenic lines in the zebrafish *bncr*^*-/-*^ background, we tested whether the amino acids in these sites alone or in combination were sufficient to switch the specificity of one species’ Bncr to favor the other species’ sperm by substituting the residues from one species’ Bncr to that of the other and vice versa. Introduction of zebrafish amino acids into medaka Bncr increased compatibility with zebrafish sperm *in vivo* and *in vitro*, particularly when introduced in combination (Fig. 5C; Suppl. Fig. 5A). In contrast, none of the tested medaka amino acid substitutions in zebrafish Bncr were sufficient to enable fertilization by medaka sperm, and neither did they disrupt compatibility with zebrafish sperm (Fig. 5D; Suppl. Fig. 5B-C). Substituting both A45 and L63 for R in medaka Bncr was sufficient to cause a clear shift in bias toward zebrafish sperm, suggesting that the positively charged patch mediated by these arginine residues is beneficial for zebrafish sperm interaction (Fig. 5E). Because medaka sperm retained compatibility with all of the tested medaka Bncr substitution mutants, other features within Bncr are required to determine medaka specificity (Fig. 5C).

### Medaka Bouncer requires N-glycosylation in finger 2

To uncover these features, we examined all tested sequences (fish Bncr homologs, ancestral states, and medaka/zebrafish Bncr chimeras) for elements that are always present when compatible with one species’ sperm but not both. All constructs that can rescue fertilization with medaka sperm contain a single predicted N-glycosylation site in finger 2 (NXS/T, where X is any amino acid except proline), while zebrafish and carp Bncr contain N-glycosylation sites in fingers 1 and 3 (Fig. 4A). In line with a previous observation that non-glycosylated zebrafish Bncr is functional with zebrafish sperm (*10*), we hypothesized that the presence of N-glycosylation in finger 2 may contribute to the medaka-specific requirement for compatibility that is not shared with the zebrafish Bncr interaction partner and manifests as asymmetrical specificity. To test the role of both number and position of Bncr N-glycosylation sites in medaka/zebrafish specificity, we generated transgenic lines in the zebrafish *bncr*^*-/-*^ background expressing medaka and zebrafish Bncr N-glycosylation site variants that we confirmed by western blotting to exhibit the expected N-glycosylation patterns (Suppl. Fig. 5D; Suppl. Data File 3).

We found that any changes to the N-glycosylation pattern of zebrafish Bncr, even when mimicking the N-glycosylation pattern of medaka Bncr with only finger 2 glycosylated, maintained zebrafish sperm compatibility and were not sufficient to rescue medaka sperm compatibility (Fig. 5F, left; Suppl. Fig. 5E). In contrast, medaka Bncr N-glycosylation variants revealed a strict requirement for finger 2 N-glycosylation: removal of this N-glycosylation site abrogated fertilization with both medaka and zebrafish sperm (Fig. 5F, right; Suppl. Fig. 5F). Rescue with either sperm could not be restored by adding an N-glycosylation site to medaka Bncr on finger 1, 3, or both despite membrane expression of all constructs (Fig. 5F, right; Suppl. Fig. 5G). Thus, this data supports finger 2 N-glycosylation in medaka Bncr as a medaka-specific requirement for sperm compatibility or protein function that is absent in medaka-incompatible Bncr proteins. However, while necessary, this N-linked glycan is not sufficient to enable medaka sperm compatibility with zebrafish Bncr.

## Discussion

In this study we investigated the role of Bncr in medaka and its features that determine species-specificity between medaka and zebrafish sperm. Examination of the medaka *bncr* locus revealed the presence of two splice isoforms, Bncra and Bncrb, that are present in many fish species (Suppl. Fig. 1A). These splice isoforms likely arose by gene duplication, which has been shown to influence the evolution of fertilization proteins in many species (reviewed in (*33*)). In the case of zebrafish, however, Bncrb appears to have been lost. Characterization of Bncra and Bncrb in medaka revealed that while Bncra is required for fertilization like zebrafish Bncr, neither male nor female medaka Bncrb mutants had any apparent fertilization defects and transgenic medaka Bncrb failed to rescue fertilization with medaka sperm when expressed in zebrafish eggs (Fig. 1D; Suppl. Fig. 1C). Bncra (Bncr) is therefore conserved as an essential fertilization factor in distantly related fish species, but the precise role of Bncrb in the egg requires further study. Bncrb may support other fertilization proteins, for example in sperm chemoattraction to the egg, but it is not necessary for this process.

Our investigation into Bncr’s role in mediating species-specific fertilization in fish revealed that instead of exhibiting strong selectivity for conspecific sperm as observed for zebrafish and medaka ((*10*) and this study), Bncr homologs in general maintain more widespread compatibility among species than previously expected (Fig. 2). Indeed, the strong specificity between medaka and zebrafish sperm-Bncr pairs appears restricted to these two species but may extend to gamete interactions between fish from genus *Oryzias* vs. suborder *Cyprinoidei* in general given the incompatibility between medaka sperm and carp Bncr.

Two important themes emerge from these observations. First, species-specific Bncr-sperm interaction can be partially overcome by expression level, reminiscent of the concentration-dependent interactions previously seen with lysin and Bindin (*17, 34*). Secondly, zebrafish sperm interact more indiscriminately with the tested Bncr homologs compared to medaka sperm, exhibiting a wider range of compatibility. This promiscuity may contribute to the observed high frequency of hybridization among species within *Cyprinidae* (*35*) and may in part explain the ability of *Danio* species to hybridize with one another and other species within *Cyprinidae* (*36*). This is further underscored by the fact that unlike previously described species-specific fertilization factor pairs like Bindin/EBR1 (*37, 38*) and lysin/VERL (*39, 40*), Bncr’s evolution is marked mostly by negative rather than positive selection. By maintaining cross-species compatibility, Bncr may have played a part in allowing cross-fertilization and hybridization of diverse fish species, particularly those without other modes of reproductive isolation. Bncr’s mammalian homolog, SPACA4, is expressed on sperm and is required for ZP binding and penetration (*8*), but whether it confers species specificity to this process has yet to be investigated.

The features in Bncr that dictate medaka or zebrafish compatibility are not mutually exclusive and comprise a different set of requirements involving a combination of specific amino acids and N-glycosylation pattern for interaction with each species’ sperm. As shown for fugu and seahorse Bncr proteins, a Bncr protein can fulfill the requirements for interaction with both medaka and zebrafish sperm simultaneously. Constituent amino acids in finger 3 appear critical for maintaining successful sperm interaction for both zebrafish and medaka, but the context is decisive (Fig. 3B-C). Specifically, introducing L63 (in zebrafish R63) into finger 3 in a medaka Bncr-like context contributes to a clear preference for medaka over zebrafish sperm as revealed by comparing fertilization rates for ancestral state nodes E and F (Fig. 4C-D). Introducing L63 into an otherwise zebrafish Bncr protein, however, is not sufficient to disrupt zebrafish compatibility nor enable medaka compatibility (Fig. 5D). When both A45 and L63 in medaka Bncr are mutated to R as in zebrafish Bncr, this mutant shifts in bias toward zebrafish sperm, suggesting that the positively charged surface provided by these residues is beneficial for zebrafish sperm, yet medaka sperm are not deterred from interaction (Fig. 5E).

While finger 3 is important for interaction with both species’ sperm, our analysis revealed that N-glycosylation is an additional context-dependent feature with differential influence on medaka and zebrafish sperm interaction. The still unknown Bncr interaction partner in zebrafish tolerates both a lack of N-glycosylation in fingers 1 and 3 (*10*) and the presence of N-glycosylation in finger 2 (Fig. 5F) in zebrafish Bncr. In contrast, removal of N-glycosylation from finger 2 of medaka Bncr disrupts function of the protein with either sperm and cannot be rescued by addition of N-glycosylation to finger 1, 3, or both (Fig. 5F). This may be a result of failed protein folding or trafficking to the membrane, yet all medaka Bncr N-glycosylation variants were still detected at the egg membrane in transgenic zebrafish lines (Suppl. Fig. 5G). Importantly, all medaka-compatible Bncr sequences have finger 2 N-glycosylation (Fig. 3B-C and 4A), giving credence to the idea that this feature is required for medaka sperm compatibility but is not sufficient alone (Fig. 5F). We therefore propose that finger 2 N-glycosylation is necessary for medaka sperm-Bncr interaction but additional features within finger 3 are also required.

To date, only three sperm proteins (Dcst1, Dcst2, and Spaca6) have been reported as essential for fertilization in zebrafish (*4, 6*), yet none of them have been shown to act as Bncr’s interaction partner. Although the identity of Bncr’s interaction partner remains elusive, this study provides valuable insights into the amino acid sites and protein features within Bncr that are needed for binding sperm, thereby shedding light on what is required by the unknown interaction partner. To reconcile the observations that the medaka and zebrafish Bncr interaction partners exhibit asymmetrical specificity yet can interact with the fugu Bncr protein with comparable efficiency, we propose three possible explanations. Either the medaka and zebrafish interaction partners are entirely different molecules, or the binding site for the two species’ interaction partners on Bncr is different with unequal binding affinities. Alternatively, the zebrafish Bncr interaction partner on sperm may have an overall higher expression that results in higher avidity even when presented with a suboptimal Bncr with lower affinity. Such a strategy would ensure efficient sperm binding to the egg with risk of binding to heterospecific eggs, which is consistent with the ability of zebrafish to hybridize with many other fish species. Identification of Bncr’s interaction partner(s) on sperm will enable differentiating between these possibilities to reveal the molecular nature of Bncr’s essential lock-and-key mechanism.

## Materials and Methods

### Ethics statement

All animal experiments were conducted according to Austrian and European guidelines for animal research and approved by the Amt der Wiener Landesregierung, Magistratsabteilung 58 - Wasserrecht (animal protocols GZ 342445/2016/12 and MA 58-221180-2021-16 for work with zebrafish; animal protocol GZ: 198603/2018/14 for work with medaka).

### Zebrafish and medaka husbandry

Zebrafish (*Danio rerio*) were raised according to standard protocols (28°C water temperature; 14/10 hour light/dark cycle). TLAB fish, generated by crossing zebrafish AB with stocks of the natural variant TL (Tüpfel long fin), served as wild-type zebrafish for all experiments. Wild-type medaka (*Oryzias latipes*, CAB strain) were raised according to standard protocols (28°C water temperature; 14/10 hour light/dark cycle) and served as wild-type medaka. *Oryzias curvinotus* were raised under the same conditions. *Bouncer* mutant zebrafish and medaka Bouncer-expressing transgenic zebrafish lines have been published previously (*10*).

### Generation of medaka *bncra* and *bncrb* mutants

Medaka *bncra* and *bncrb* mutants were generated using Cas9-mediated mutagenesis. Guide RNAs (sgRNAs) targeting the third (*bncra*) and second exons (*bncrb*) (see table below) were synthesized by *in vitro* transcription using the MEGAscript T7 Transcription Kit (ThermoFisher) after annealing oligos according to (*41*).

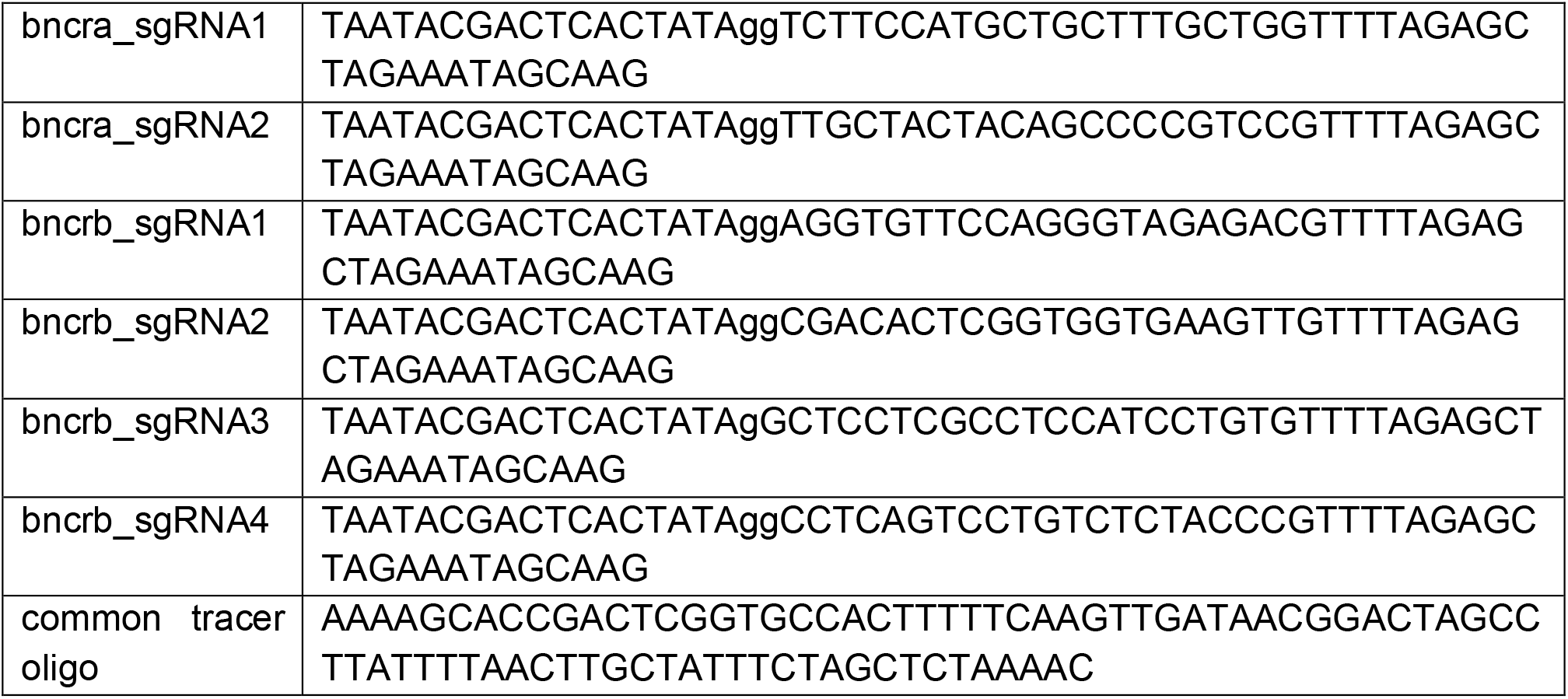

*Cas9* mRNA was synthesized using the mMESSAGE mMACHINE SP6 Transcription Kit (ThermoFisher) using a linearized pCS2 vector template containing the Cas9 ORF (*41*). One-cell medaka embryos (CAB strain) were co-injected with *cas9* mRNA and sgRNAs in 1X Yamamoto’s ringer’s solution (1.00 g NaCl, 0.03 g KCl, 0.04 g CaCl_2_·2H_2_O, 0.10 g MgCl_2_·6H_2_O, 0.20 g NaHCO_3_ in 1000 mL, pH 7.3). Potential founder fish were crossed to wild-type CAB fish; the offspring from these crosses were screened by PCR for mutations in *bncra* (bncra_F: AGTACAAGCATCTGAGTAGGG and bncra_R: AGGCTGTGAACCTGACTG) and *bncrb* (bncrb_F: AGAGGCCTTTATAATGTGGACA and bncrb_R: CCATCTCATAGGAACCACAGA) based on a shift in amplicon size compared to wild-type. Offspring of founder fish were raised to adulthood and in-crossed to produce homozygous mutants. The 5-nt and 38-nt deletions in exons 3 and 2, respectively, were detected by PCR and confirmed by Sanger sequencing to be frameshift mutations in *bncra* and *bncrb*, respectively. The wild-type and mutant sequences and corresponding translated amino acid sequences are provided in Suppl. Data File 1. Genotyping of *bncra* and *bncrb* mutants was done using PCR with the primers given above and standard gel electrophoresis using a 4% agarose gel.

### Quantification of medaka *in vivo* fertilization rates

To quantify fertilization rates of wild-type and mutant medaka, mating crosses were set up the night before inside the tanks in the fish water system. One male per two or three females was set up in the same tank; the male and females were separated with a vertical divider which was removed the morning of egg collection. After mating, the eggs were collected carefully with a fine mesh net using the thumb and forefinger to remove them from each female’s body and placed into a separate petri dish containing 1X Yamamoto’s ringer’s solution (1.00 g NaCl, 0.03 g KCl, 0.04 g CaCl_2_·2H_2_O, 0.10 g MgCl_2_·6H_2_O, 0.20 g NaHCO_3_ in 1000 mL, pH 7.3). The eggs from each female were visually inspected under a dissection microscope and the number of unactivated eggs was recorded. These are easily distinguished from activated eggs based on their dark appearance, higher density of cortical alveoli, and the close apposition of the chorion to the egg membrane. Approximately 2-3 hours post-collection, the eggs were inspected again, and fertilization rates were quantified based on the presence of cell cleavage.

### Generation of transgenic zebrafish lines

All zebrafish transgenic lines were generated using Tol2-mediated transgenesis. To generate plasmids encoding other fish Bncrs, chimeras, and ancestral states for transgenesis, each Bncr ORF lacking its endogenous signal peptide sequence but including the C-terminal tail was ordered as a custom gBlock (IDT) and Gibson cloned into a vector containing Tol2 sites, the *actb2* promoter, and the zebrafish Bncr signal peptide followed by sfGFP. Each Bncr sequence was inserted in frame downstream of sfGFP such that the resulting plasmids were as follows: Tol2 – *actb2* promoter – zebrafish Bncr SP – sfGFP – Bncr sequence plus C-terminal tail – SV40 UTR – Tol2. All amino acid substitution constructs were generated using PCR-based site-directed mutagenesis of plasmids containing the wild-type medaka or zebrafish Bncr ORF as the template. All transgene sequences are provided in Suppl. Data File 3. To generate transgenic zebrafish lines, *tol2* mRNA was co-injected with the plasmid encoding the desired Bncr transgene into one-cell stage zebrafish embryos from a ♀ *bncr*^+/-^ x ♂ *bncr*^−/-^ cross. Larvae were screened for fluorescence one day post-fertilization (1 dpf) and grown to adulthood. Potential founders were crossed to *bncr*^+/-^ or *bncr*^−/-^ zebrafish and their progeny (F1) was grown to adulthood if fluorescent at 1 dpf. Homozygous *bncr* mutant F1 and F2 fish stably expressing the desired transgene were used for experimentation.

### Transgenic egg imaging with CellMask

Transgenic zebrafish females were set up with males as previously described. On the morning of collection, females were either allowed to mate with males or were squeezed according to the IVF protocol. Eggs were collected immediately in blue water (3 g Instant Ocean® sea salt per 10 L fish system water, 0.0001% (w/v) methylene blue) and allowed to activate for ∼10 minutes. As soon as their chorions were lifted, 15-20 eggs were manually de-chorionated with fine forceps in a Silguard dish filled with 1X Danieau’s solution. Eggs were incubated at RT with gentle rocking for 15 minutes in a watch glass containing 0.01 mg/ml CellMask Deep Red plasma membrane stain (Invitrogen) in 250 µl 1X Danieau’s solution and covered with aluminum foil. After incubation, the eggs were transferred to a new watch glass containing 1X Danieau’s solution and were then imaged immediately in an agarose mold filled with 1X Danieau’s solution using an upright point laser scanning confocal microscope (LSM800 Examiner Z1, Zeiss) with a 10X/0.3 N-achroplan water objective.

### Quantification of *bncr* transgene copy number by qPCR

Primers targeting medaka *bncra* (medbncr_F: TCAGGTTCACAGCCTACGTC and medbncr_R: GTTACAGTACGGCCAGTCACA) and zebrafish *bncr* (zfbncr_F: CACCAGATGATCCGGGGAAA and zfbncr_R: CTGGGAGTTGCAGTAGTGTCC) were directly compared for efficiency by amplification of a dilution series of a template plasmid containing one copy of each transgene. The template plasmid was cloned for this purpose using a pBluescript II SK(+) vector (gift from Katharina Lust, Tanaka lab) and contained the following elements: I-SceI site – medaka *actb* promoter – zebrafish Bncr SP – mCherry – mature zebrafish Bncr plus C-terminal tail – SV40 UTR – mature medaka Bncra plus C-terminal tail – I-SceI site. Linear regression analysis of the standard curve yielded a line of best fit; the equation of which could be used to calculate copy number based on Cq value for each primer pair. Eggs from three medaka or zebrafish females per line were collected immediately after laying and homogenized in TRIzol (ThermoFisher Scientific). Total RNA from each egg sample was isolated by standard phenol/chloroform extraction. cDNA synthesis was done using the iScript cDNA Synthesis Kit (Bio-Rad); for each sample, 500 ng of input RNA was used. qPCR was performed in technical duplicates using 2X GoTaq Master Mix (Promega) and the primers given above depending on the target transgene. Copy number was then calculated based on the average Cq value of technical duplicates for each biological replicate using the equation derived for each primer pair.

### *In vitro* fertilization assay with medaka and zebrafish

To collect zebrafish eggs and sperm for *in vitro* fertilization, wild-type TLAB zebrafish males were set up the night before experimentation with transgenic or wild-type zebrafish females in a small, plastic breeding tank with a divider separating the two fish. On the day of experimentation, sperm was collected from zebrafish males after anesthetization in 0.1% (w/v) tricaine (25X stock solution in dH_2_O, buffered to pH 7-7.5 with 1 M Tris pH 9.0) in fish system water. Using plastic tubing with a capillary in one end and a pipette filter tip in the other end, sperm was mouth-pipetted from the urogenital opening of each male placed belly-up in a slit in a sponge wetted with fish water. Sperm was transferred directly to a 1.5-mL tube containing Hank’s saline (0.137 M NaCl, 5.4 mM KCl, 0.25 mM Na_2_HPO_4_, 0.44 mM KH_2_PO_4_, 1.3 mM CaCl_2_, 1.0 mM MgSO_4_, 4.2 mM NaHCO_3_) on ice. To collect zebrafish eggs, transgenic or wild-type zebrafish females were anesthetized on the day of experimentation as described above. After anesthetization, the abdomen of each female was carefully dried on a paper towel and the fish was transferred to a petri dish. Gentle pressure was applied to the belly of the female with the thumb of one hand while her back was supported with a finger of the other hand. Approximately 50-100 eggs were exuded from the female before transferring her to a second petri dish and repeating the process. The fish was immediately placed back into fish water, and one clutch of eggs was fertilized with zebrafish sperm, and the other with medaka sperm, such that the same number of sperm were used on both clutches of eggs. 500 µl of blue water was added immediately to each clutch after sperm addition to activate gametes. The dishes were left undisturbed for 3-5 minutes and then filled with blue water and placed into an incubator at 28°C.

Male medaka were kept in a tank without females at least one week prior to sperm collection. Medaka sperm was collected from medaka males in the same manner as for zebrafish. For generation of unfertilized, unactivated medaka eggs, infertile hybrid *O. curvinotus* x *O. latipes* males were used to mate with wild-type CAB females. These crosses were set up the night before experimentation inside their tanks in the fish water system with a vertical divider separating one male from two to four females or two males from four to five females. Freshly spawned medaka eggs were collected directly from the bodies of females using a net with fine mesh and by gently pulling the eggs from the fish in the net using the thumb and forefinger on the outside of the net, taking care not to crush any eggs during removal. Collected eggs were placed into petri dishes containing 1X Yamamoto’s ringer’s solution. After collection, eggs were visually inspected under a dissection microscope to separate them and remove any crushed or activated eggs and were then divided into two separate dishes. As much ringer’s solution as possible was removed from each dish such that the eggs remained submerged when the dish was tilted on its lid. The volume of each species’ sperm suspension needed to have medaka and zebrafish sperm in equal numbers was pipetted directly onto the eggs in each dish. For fertilization with medaka sperm, 500 µl of blue water was added immediately after sperm addition to activate the sperm (medaka sperm are not as active in ringer’s solution, but zebrafish sperm are). For fertilization with zebrafish sperm, 2 minutes after sperm addition, 2 µL of 0.1% (w/v) calcimycin in DMSO was pipetted carefully onto the eggs to activate them (*26*). After 10 minutes, the dishes were filled with blue water before being placed into an incubator at 28°C.

In general, based on the number of egg clutches of eggs to be fertilized in each experiment, one male was used per 100 µL of Hank’s saline. Because sperm is used in great excess during IVF, any concentration above 50,000 sperm/µl was used. Sperm were counted manually in a Neubauer chamber to ensure that the same number of medaka and zebrafish sperm was used on each sample in the same experiment. In general, 3-4 million sperm were used to fertilize each clutch of eggs. Fertilization rates were quantified approximately 3 hours after IVF by using a dissection microscope and counting the number of fertilized embryos with cell cleavage and unfertilized eggs that had remained at the one-cell stage and did not develop.

### Positive selection analysis

Mature Bncr protein sequences were aligned using MAFFT (*42*) and codon alignment was generated using PAL2NAL (*43*). Codon alignments were then used as input into IQ-TREE (*44*) to generate the best substitution model and a maximum-likelihood tree was generated using 1000 ultrafast bootstrap iterations (*45*). Codon alignments and the maximum-likelihood tree were used as input into HyPhy (*46*) to test the mode of selection acting on Bncr in fish. A suite of tests was performed across all sequences by using MEME (*47*), FUBAR (*48*), FEL (*49*) and BUSTED (*50*). The mode of selection acting on the zebrafish and medaka lineages was tested by selecting on the branches leading to these lineages and performing aBSREL (Smith et al., 2015) and Contrast-FEL (*51*). We further mapped the residues identified to be under selection regimes of interest onto the predicted 3-D structures of zebrafish and medaka Bncr. In addition, the level of conservation was mapped onto the 3-D structures for zebrafish and medaka Bncr using CONSURF (*52*) and visualized using PyMOL (http://www.pymol.org).

### Prediction of Bncr ancestral states

Bncr amino acid sequences were aligned using MUSCLE (*53*) with default parameters. A phylogeny was then reconstructed using MrBayes and mcmc=2000000 (*54*)). Finally, ancestral amino acid states were reconstructed for all nodes of the obtained phylogeny with PAML with default parameters (*55*). The alignment, phylogeny, and ancestral reconstructions as well as the relevant control files are available on GitHub (https://github.com/kristabriedis/AncestralBncrs).

### De-glycosylation and western blot analysis

To collect egg cap lysates for de-glycosylation enzyme treatment and western blot analysis, transgenic zebrafish females for each line of interest and a male were set up the night before in a mating tank and separated with a plastic divider. On the morning of collection, the fish were allowed to mate, and their eggs were collected immediately in blue water. As soon as the chorions were lifted, the eggs were de-chorionated and de-yolked manually with fine forceps in a Silguard dish filled with 1X Danieau’s solution (58 mM NaCl, 0.7 mM KCl, 0.4 mM MgSO_4_, 0.6 mM Ca(NO_3_)_2_, 5 mM HEPES, pH 7.6). Eggs were dissected 5 at a time such that a total of 20 egg caps were collected in 8 µl of 1X Danieau’s solution and were immediately pipetted into a tube on dry ice. Samples were kept at −70°C until processing. To each sample, 32 µl of nuclease-free water was added and samples were divided equally into two tubes. The untreated sample was kept at −70°C, while the other was treated overnight with Protein Deglycosylation Mix II (NEB) using non-denaturing reaction conditions according to the manufacturer’s protocol. All samples were boiled with 1X Laemmli buffer containing 10% ß-mercaptoethanol before SDS-PAGE using Mini-PROTEAN TGX (Bio-Rad) pre-cast gels. After SDS-PAGE, samples were wet-transferred onto a nitrocellulose membrane which was blocked with 5% milk powder in 0.1% Tween-20 in 1X TBS (TBST). Membranes were incubated in primary rabbit anti-GFP antibody [1:1000, (A11122, Invitrogen)] overnight at 4°C, then washed with TBST before HRP-conjugated secondary antibody [1:10.000 (115–036–062, Dianova)] incubation for 30 min to 1 hr. Membranes were washed a few times in TBST before HRP activity was visualized using Clarity Western ECL Substrate (Bio-Rad) on a ChemiDoc (Bio-Rad). For visualizing tubulin levels, membranes were stripped using Restore Western Blot Stripping Buffer (ThermoFisher Scientific) before washing, blocking, and incubation with mouse anti-alpha-tubulin antibody [1:20.000 (T6074, Merck)] and proceeding with secondary antibody staining and detection as described above.

### Statistical analysis

Statistical comparisons between medaka *bncra* and *bncrb* mutants vs. wild-type and the transgenic *bncra* rescue line were performed using the Kruskal-Wallis test with Dunn’s multiple comparisons test at the 0.95 confidence level. Statistical comparisons between clutches of eggs fertilized by medaka vs. zebrafish sperm in IVF experiments were made using the Wilcoxon matched-pairs signed rank test with method of Pratt, such that pairs for which the two values were equal were not excluded. The median of the bias indices derived from paired IVF data was tested for being significantly different from a hypothetical median of 0 (indicating no bias) using the Wilcoxon Signed Rank Test at the 0.95 confidence level. The method of Pratt was used such that median values equal to 0 were not excluded.

## Acknowledgements

We thank Manfred Schartl for sharing RNA-seq data from medaka ovaries and testes prior to publication; Maria Novatchkova for help with RNA-seq analysis; Katharina Lust for advice on medaka techniques; Luca Jovine for valuable insight, fruitful discussions, and help with structural predictions; Felicia Spitzer, Mirjam Binner, and Anna Bandura for help with genotyping; Friedrich Puhl, Julia Koenig, and Dijana Sunjic for taking care of zebrafish and medaka; and the Pauli lab for helpful discussions about the project and feedback on the manuscript. KRBG was supported by a DOC Fellowship from the Austrian Academy of Sciences. Work in the Pauli lab was supported by the FWF START program (Y 1031-B28 to AP), the HFSP Career Development Award (CDA00066/2015 to AP), a HFSP Young Investigator Award (RGY0079/2020 to AP) and the FWF SFB RNA-Deco (project number F80). The IMP receives institutional funding from Boehringer Ingelheim and the Austrian Research Promotion Agency (Headquarter grant FFG-852936). Work by JMS and YM in this project was supported by the Israel Science Foundation grant 636/21 to YM. For the purpose of Open Access, the author has applied a CC BY public copyright license to any Author Accepted Manuscript (AAM) version arising from this submission.

## Data and Materials Availability

All data needed to evaluate the conclusions in the paper are presented in the paper and/or the Supplementary Materials. Files pertaining to ancestral state reconstruction are available on GitHub (https://github.com/kristabriedis/AncestralBncrs).

## Author Contributions

KRBG and AP conceived the study; KRBG designed, performed, and analyzed experiments, with assistance from KP in some IVF experiments, *in vivo* fertilization rate collection, and western blotting, and BSS in some IVF experiments. AS conducted phylogenetic analysis. JS and YM performed positive selection analysis. FK generated predicted ancestral states of Bouncer. AP supervised the study. KRBG wrote the original draft. KRBG and AP revised the manuscript with input from all authors.

## Supplementary Materials for

**Fig. S1.**
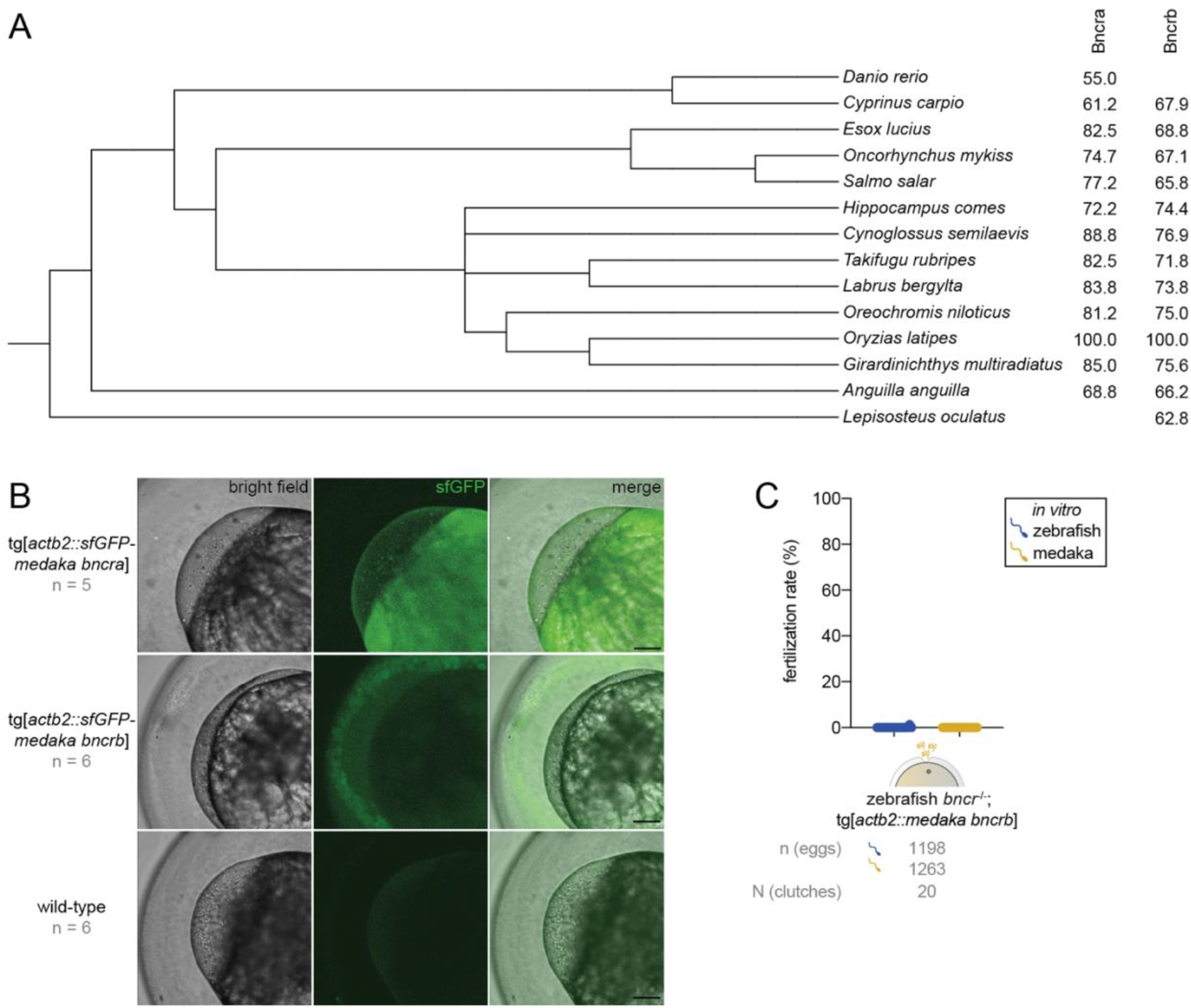
**(A)** Taxonomic tree depicting the presence/absence of Bncra and Bncrb in selected fish species. All Bncr homologs of the respective species were assigned to either Bncra or Bncrb subfamilies based on the highest similarity to its medaka ortholog. **(B)** Confocal maximum intensity Z-projections of transgenic zebrafish *bncr*^*-/*-^ eggs expressing sfGFP-tagged medaka Bncra (top) and Bncrb (middle). Wild-type zebrafish eggs with no transgene are shown below. All images are taken at 10X magnification. Scale bar = 100 µm. **(C)** Medaka/zebrafish IVF with transgenic zebrafish *bncr*^*-/*-^ eggs expressing sfGFP-tagged medaka Bncrb.

**Fig. S2.**
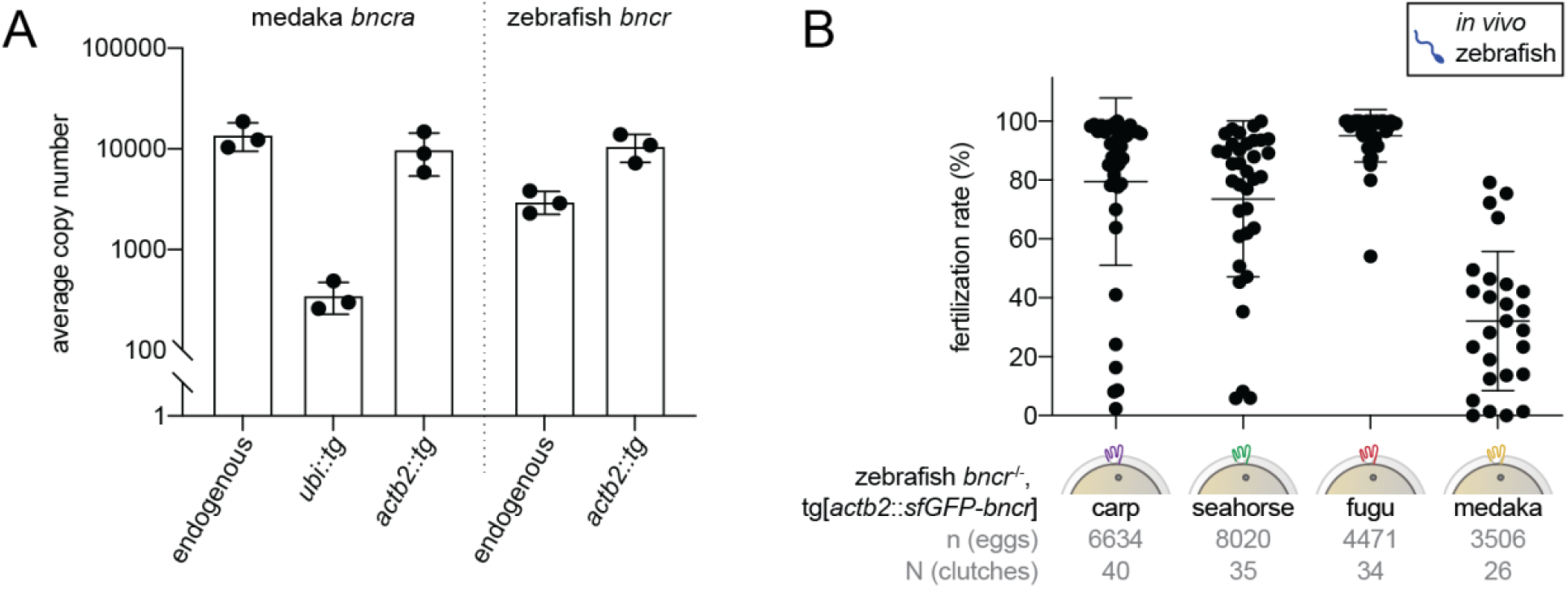
**(A)** Average copy number (from 3 biological replicates) of endogenous, *ubiquitin* promoter-driven, and *actin* promoter-driven medaka *bncra* in wild-type medaka eggs and transgenic zebrafish *bncr*^*-/-*^ eggs as measured by qPCR (left). Average endogenous and *actin* promoter-driven copy numbers for zebrafish *bncr* in wild-type and transgenic zebrafish *bncr*^*-/-*^ eggs, respectively, as measured by qPCR (right). Y-axis is plotted in log_10_ scale. **(B)** *In vivo* fertilization rates of transgenic zebrafish *bncr*^*-/-*^ lines expressing carp, seahorse, fugu, and medaka *bncra*. Fertilization rates with an *actin* promoter-driven zebrafish *bncr* rescue line were previously reported (Herberg et al., 2018). Because each line may have a different expression level of its respective transgene vs. another, statistical comparisons between lines were not performed.

**Fig. S3.**
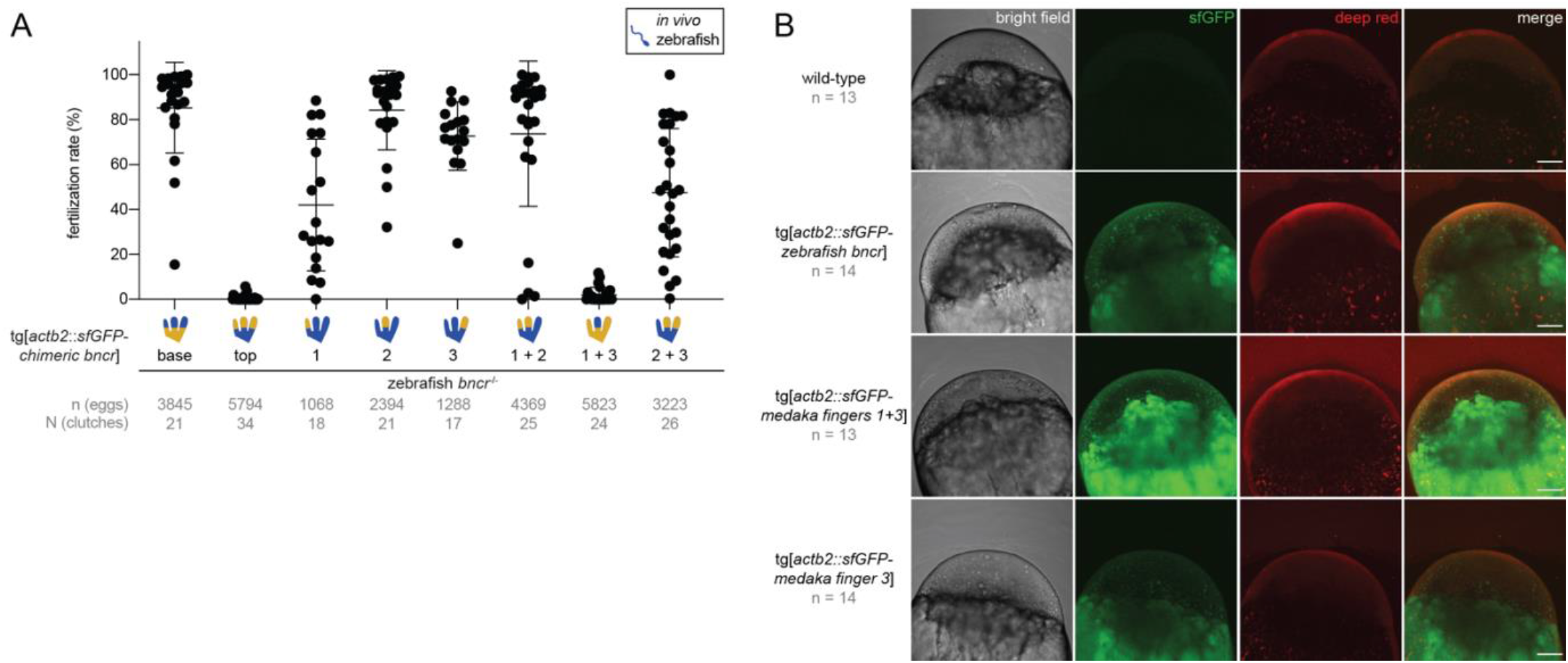
**(A)** *In vivo* fertilization rates of transgenic zebrafish *bncr*^*-/-*^ lines expressing sfGFP-tagged chimeric medaka/zebrafish Bncr constructs. The “finger(s)” or region(s) of Bncr that were changed from the zebrafish sequence to that of medaka are indicated below the X-axis. Because each line may have a different expression level of its respective transgene vs. another, statistical comparisons between lines were not performed. **(B)** Confocal maximum intensity Z-projections of wild-type (top) and transgenic zebrafish *bncr*^*-/*-^ eggs (below). Chimeric sfGFP-tagged Bncr constructs medaka fingers 1 + 3 (3^rd^ row) and medaka finger 3 (bottom row) show expression at the egg membrane, similar to zebrafish *bncr* (2^nd^ row). Red, CellMask Deep Red membrane stain. All images are taken at 10X magnification. Scale bar = 100 µm.

**Fig. S4.**
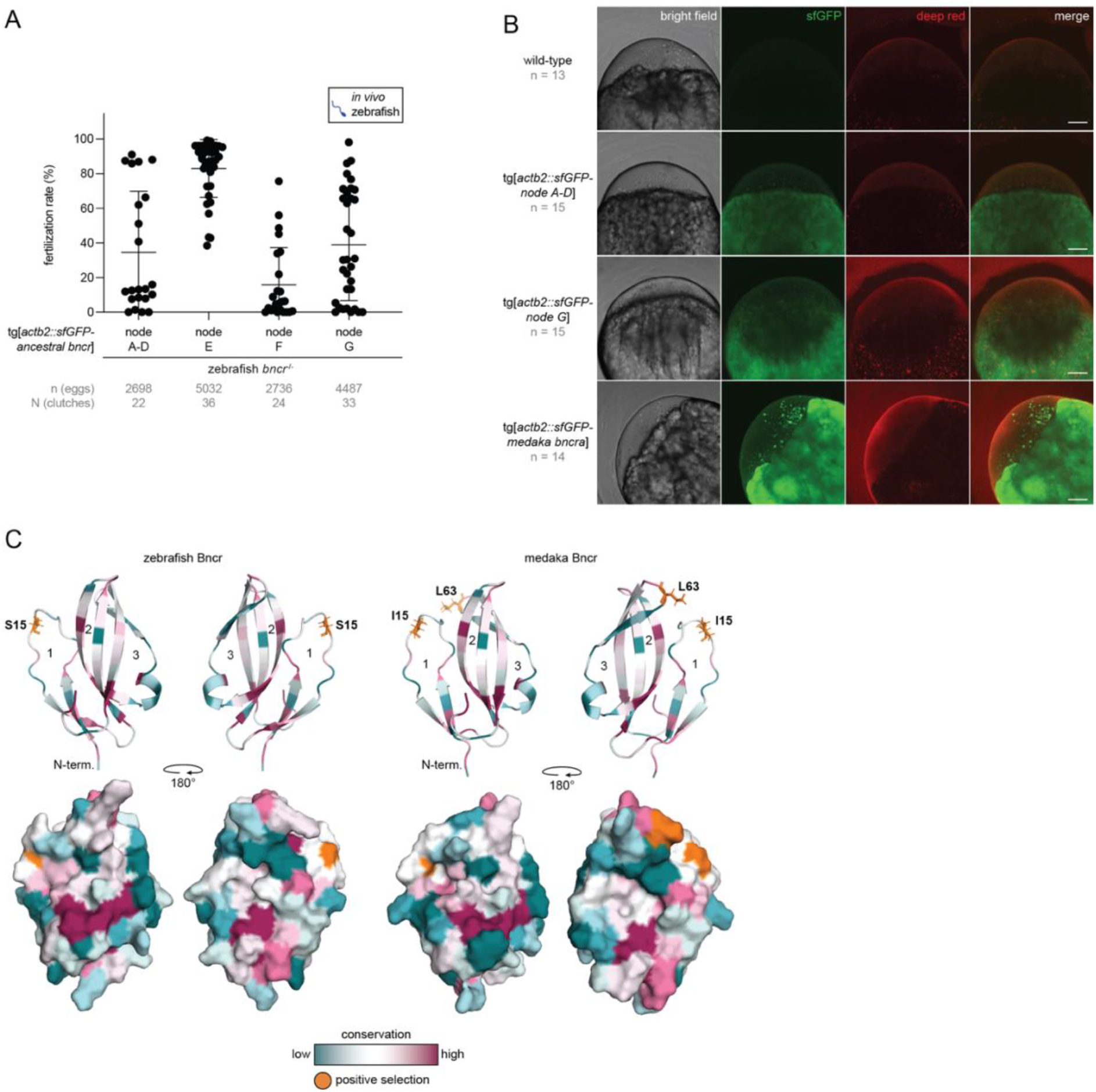
**(A)** *In vivo* fertilization rates of transgenic zebrafish *bncr*^*-/-*^ lines expressing sfGFP-tagged ancestral Bncr states at nodes A-D, E, F, and G. Because each line may have a different expression level of its respective transgene vs. another, statistical comparisons between lines were not performed. **(B)** Confocal maximum intensity Z-projections of wild-type (top) and transgenic zebrafish *bncr*^*-/*-^ eggs (below). Ancestral sfGFP-tagged Bncr constructs nodes A-D (2^nd^ row) and node G (3^rd^ row) show expression at the egg membrane, though are more weakly expressed than medaka *bncra* (bottom row). Red, CellMask Deep Red membrane stain. All images are taken at 10X magnification. Scale bar = 100 µm. **(C)** Cartoon and surface representation models of zebrafish (left) and medaka (right) Bncr proteins with sites colored according to conservation level. Site 15 is under positive selection in both zebrafish and medaka Bncr, whereas site 63 is under positive selection specifically in the medaka lineage. Amino acids in positively selected sites are colored orange; conservation level ranges from low (dark teal) to high (dark magenta).

**Fig. S5.**
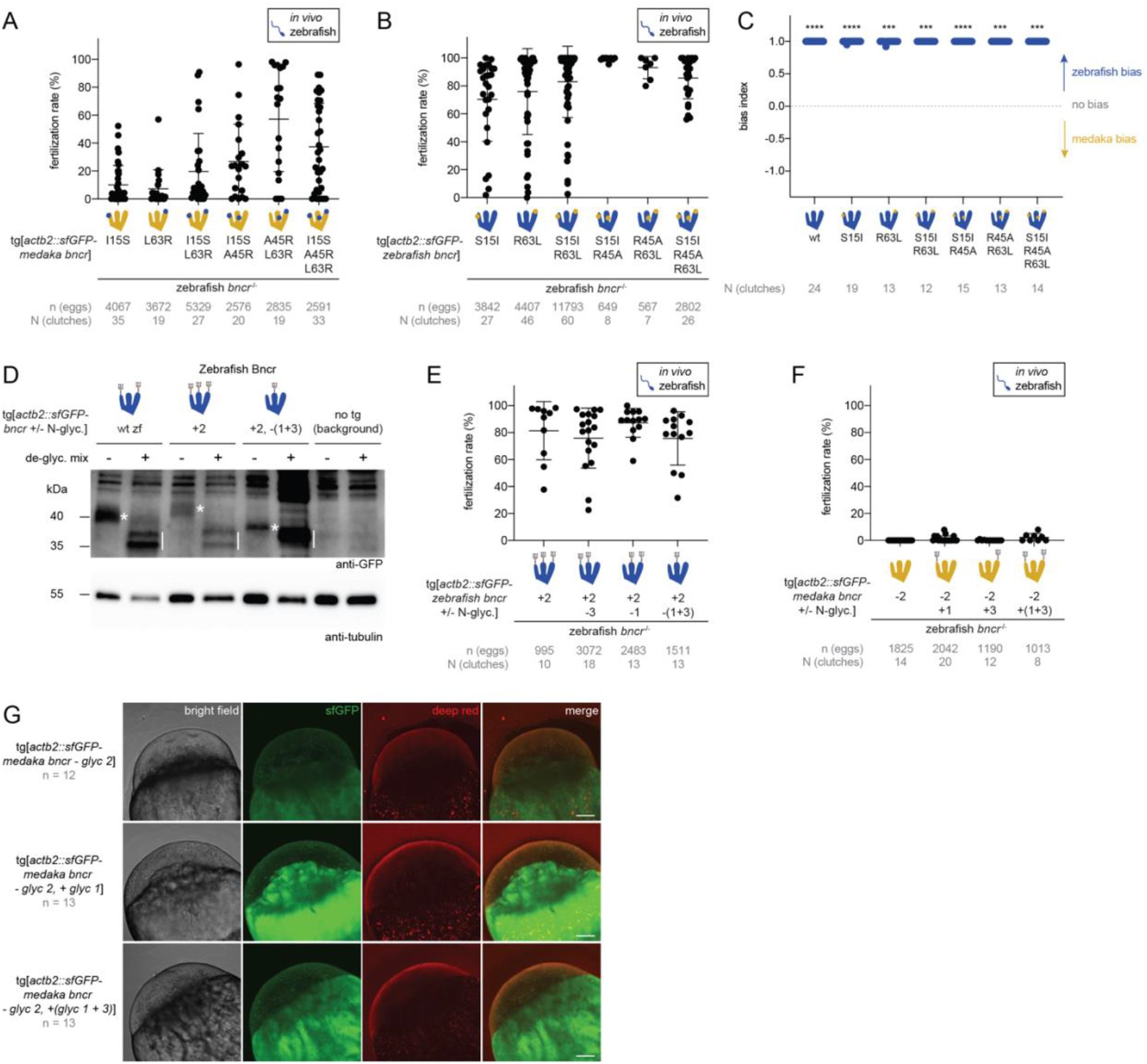
**(A)** *In vivo* fertilization rates of transgenic zebrafish *bncr*^*-/-*^ lines expressing sfGFP-tagged medaka Bncr constructs with zebrafish amino acid substitutions. **(B)** *In vivo* fertilization rates of transgenic zebrafish *bncr*^*-/-*^ lines expressing sfGFP-tagged zebrafish Bncr constructs with medaka amino acid substitutions. **(C)** Bias index derived from IVF data with zebrafish Bncr amino acid substitution constructs from Figure 5B. All constructs show bias for zebrafish sperm. (Wilcoxon signed rank test vs. theoretical median of 0 with method of Pratt: ****P < 0.0001 (medaka vs. zebrafish sperm with zebrafish Bncr, zebrafish Bncr S15I, and zebrafish Bncr S15I, R45A), ***P = 0.0002 (zebrafish Bncr R63L, zebrafish Bncr R45A, R63L), ***P = 0.0005 (zebrafish Bncr S15I, R63L), ***P = 0.0001 (zebrafish Bncr S15I, R45A, R63L). **(D)** Western blot with GFP antibody of zebrafish Bncr N-glycosylation variant egg lysates, untreated vs. treated with de-glycosylation enzyme mix. A higher molecular weight (m.w.), glycosylated GFP-Bouncer-signal accompanied by a smear is visible in the untreated samples (labeled with *) which shifts downward upon de-glycosylation treatment to ∼35 kDa (labeled with |). The bands from untreated samples show the highest m.w. for zebrafish Bncr with three N-glycosylation sites (+2) above 40 kDa, followed by ∼40 kDa for two N-glycosylation sites (wt zf), and below 40 kDa for one N-glycosylation site (+2, -(1+3)), in line with all constructs being N-glycosylated as expected. A lysate from wild-type zebrafish embryos (no transgene expressed) is shown on the right, indicating background signal. **(E)** *In vivo* fertilization rates of transgenic zebrafish *bncr*^*-/-*^ lines expressing sfGFP-tagged medaka Bncr N-glycosylation variants. **(F)** *In vivo* fertilization rates of transgenic zebrafish *bncr*^*-/-*^ lines expressing sfGFP-tagged zebrafish Bncr N-glycosylation variants. **(G)** Confocal maximum intensity Z-projections of transgenic zebrafish *bncr*^*-/*-^ eggs expressing medaka N-glycosylation variants. For positive and negative controls, see Suppl. Fig. 4B. All three constructs show sfGFP expression at the membrane, though weaker signal is observed with medaka Bncr - glyc 2 (top row). Red, CellMask Deep Red membrane stain. All images are taken at 10X magnification. Scale bar = 100 µm.

## Data S1. Wild-type and mutant medaka *bncra* and *bncrb* sequences

Exons are annotated in the following color code: exon 1, exon 2, exon 3

**Wild-type *bncra* cDNA; deletion indicated in <u>bold</u>**

ATGGGATCACTGAGAACCAGGCAGCTCTTCCATGCTGCTTTGCTGTGGCTTTGCCTTC CCCTTCCTCTGCTGCTCTGTGAAAACCTGCATTGCTACTACAGCCCC**<u>GTCCT</u>**GGAGA AGGAAATAACGTTTGAACTCGTCGTGACAGAATGCCCTCCGAATGAGATGTGCTTTA AGGGGTTGGGTCGCTACGGCAACTACACTGCCCTATCAGCCAGGGGCTGCATGTTGG AGAAAGACTGCAGTCAGGTTCACAGCCTACGTCTCCTGGGCACCGTCTACACCATGA GCTACAGCTGCTGTGACTGGCCGTACTGTAACCGGGCCGTCGCCCTGGAGCCGCTCA CTGCTATGCTGGTGGCTGCTGCTGTGGTGGCCTGCAGCTTTTGTCTAACATGA

**Wild-type *bncra* cDNA translation (131 amino acids)**

MGSLRTRQLFHAALLWLCLPLPLLLCENLHCYYSPVLEKEITFELVVTECPPNEMCFKGL GRYGNYTALSARGCMLEKDCSQVHSLRLLGTVYTMSYSCCDWPYCNRAVALEPLTAM LVAAAVVACSFCLT*

**Mutant *bncra* (5-nt deletion) cDNA**

ATGGGATCACTGAGAACCAGGCAGCTCTTCCATGCTGCTTTGCTGTGGCTTTGCCTTC CCCTTCCTCTGCTGCTCTGTGAAAACCTGCATTGCTACTACAGCCCCGGAGAAGGAA ATAACGTTTGA

**Mutant *bncra* (5-nt deletion) cDNA translation (41 amino acids)**

MGSLRTRQLFHAALLWLCLPLPLLLCENLHCYYSPGEGNNV*

**Wild-type *bncrb* cDNA; deletion indicated in <u>bold</u>**

ATGGGATCACTGAGAACCAGCACAATTTTGGGCCAACCTCAATTGCTCCTCGCCTCC ATCCTGTTGGTTTCTGGTCCCCTCAGTCCTGTCTCTA**<u>CCCTGGAACACCTCTTGTGT AACGTCTGCCCCCTGCAT</u>**GAAAAATCTGAGTTGTGTCCAAACTTCACCACCGAGTG TCGGCCCGGCGAGCGCTGCACCAGCTCAAGAGGCTTCTACGGTGCCCTTCACGTCCT TTCCGCTCAGGGCTGCATCAGTGCCGACCTCTGTGGTTCCTATGAGATGGTCACTTA CAGAGGAATCAAATATAAACTTCGTTATGCTTGCTGCTGCGGAAACACATGTAACGA GGCGCCTGAATCCAAAACCACACTGAAGGAGCTGCTGCAGATGATCCAAGCTAAAG CAAATGGCACTGAGGCTGCTGTGGAAAAGCCTTTGGCTGTGTGTGCAAACAACACA CTGATAGAAACCAGTGCTCCTCCTGCTGTTAAGGCATAG

**Wild-type *bncrb* cDNA translation (163 amino acids)**

MGSLRTSTILGQPQLLLASILLVSGPLSPVSTLEHLLCNVCPLHEKSELCPNFTTECRPGER CTSSRGFYGALHVLSAQGCISADLCGSYEMVTYRGIKYKLRYACCCGNTCNEAPESKTT LKELLQMIQAKANGTEAAVEKPLAVCANNTLIETSAPPAVKA*

**Mutant *bncrb* (38-nt deletion) cDNA**

ATGGGATCACTGAGAACCAGCACAATTTTGGGCCAACCTCAATTGCTCCTCGCCTCCATCCTGTTGGTTTCTGGTCCCCTCAGTCCTGTCTCTAGAAAAATCTGA

**Mutant *bncrb* (38-nt deletion) cDNA translation (34 amino acids)**

MGSLRTSTILGQPQLLLASILLVSGPLSPVSRKI*

## Data S2. Sequences in alignment for Figure 1B

>Danio_rerio_XP_005173770.1_(zebrafish Bncr) QGLRCLFCPVTSLNSSCAPVVTECPVQELCYTADGRFGRSSVLFRKGCMLRADCSRSRH QMIRGNNISFSFSCCGGHYCN

>Oryzias_latipes_H2LID1.1_(medaka Bncra) ENLHCYYSPVLEKEITFELVVTECPPNEMCFKGLGRYGNYTALSARGCMLEKDCSQVHS LRLLGTVYTMSYSCCDWPYCN

>Oryzias_latipes_H2LID5_(medaka Bncrb) EHLLCNVCPLHEKSELCPNFTTECRPGERCTSSRGFYGALHVLSAQGCISADLCGSYEMV TYRGIKYKLRYACCCGNTCN

>Cyprinus_carpio_XP_018955736.2_(carp Bncra) ENLYCYYCPQTSFNRSCRHILSECRPQELCFTALGRFGHAPVLFSKGCMSQRDCVRSSSQ MIRGNNISFTNSCCGRPYCN

>Cyprinus_carpio_XP_018955710.2_(carp Bncrb) VLLCHYCPLQAAGTRCNITTECLEHERCSSGWRRYGRVHVLALQGCLSPELCGSNQTLT HKGLEYEITYTCCCRDLCN

>Takifugu_rubripes_XP_011605859.1_(fugu Bncra) DNLLCYFSPLLEKEVSFKFIATECPPGDLCFKADGRYGNHSALSGRGCMAREACSQTHSI RYKGSVFVMSYSCCDSPYCN

>Takifugu_rubripes_XP_011605858.1_(fugu Bncrb) DTLLCYFCPLQHKTDSCVNTTSRCPPTQRCSSSRGHYGLVHVLSAQGCMDVALCGSYEI LSFKGTDFNVSHTCCCKDQCN

>Hippocampus_comes_XP_019712504.1_(seahorse Bncra) GNLRCLYRPILEKEYEFQPIVTECPRGEVCYKAEGRYGNYSALSASGCMPRRVCGLQHD L

SYQGVVYTMSYSCCDRPYCN

>Hippocampus_comes_translation from genomic frame 5KV880484.1_5 (seahorse Bncrb) TSLLCHFCPLQPKEFPCTNLTTECMPGQRCATSRAYYGVVHVLSAQGCVDARLCGNRLS VSHMGVEYRLRHSCCCKDKCN

>Salmo_salar_XP_013981439.1_(salmon Bncra) NNLLCYYSPIMYRNKTFDLILTECPPTELCMTGNGRYGNHSALSTRGCVAPTGCGQVHP LRLKGTVYTMTYACCDYNYCN

>Salmo_salar_XP_013981440.1 (salmon Bncrb)

TSLRCNFCPLQHKGRSCSNDSTTECLPQERCGTSSGRFGPIHILSAQGCLTPDLCNSTHAVTYRGVSYNVTYRCCCRDQCN

## Data S3. Transgenic line Bncr sequences

All Bncr protein sequences listed below are preceded by the zebrafish Bncr signal peptide sequence and sfGFP ORF (no stop codon) as follows:

MGCVLLFLLLVCVPVVLPTRVSKGEELFTGVVPILVELDGDVNGHKFSVRGEGEGDATNGKLTLKFICTTGKLPVPWPTLVTTLTYGVQCFSRYPDHMKQHDFFKSAMPEGYVQERTISFKDDGTYKTRAEVKFEGDTLVNRIELKGIDFKEDGNILGHKLEYNFNSHNVYITADKQKNGIKANFKIRHNVEDGSVQLADHYQQNTPIGDGPVLLPDNHYLSTQSVLSKDPNEKRDHMVLLEFVTAAGITLGMDELYKTRAAEF…

<u>Other fish Bncr homologs</u>

**Medaka Bncra**

…ENLHCYYSPVLEKEITFELVVTECPPNEMCFKGLGRYGNYTALSARGCMLEKDCSQVHSLRLLGTVYTMSYSCCDWPYCNRAVALEPFTAMLVAAAVVACSFCLT*

**Medaka Bncrb**

…LEHLLCNVCPLHEKSELCPNFTTECRPGERCTSSRGFYGALHVLSAQGCISADLCGSYEMVTYRGIKYKLRYACCCGNTCNEAPESKTTLKELLQMIQAKANGTEAAVEKPLAVCANNTLIETSAPPAVKA*

**Carp Bncra**

…ENLYCYYCPQTSFNRSCRHILSECRPQELCFTALGRFGHAPVLFSKGCMSQRDCVRSSSQMIRGNNISFTNSCCGRPYCNSSRGCDHSLALLTVSAITASVLTADWTRAGLMMPS*

**Seahorse Bncra**

…GNLRCLYRPILEKEYEFQPIVTECPRGEVCYKAEGRYGNYSALSASGCMPRRVCGLQHDLSYQGVVYTMSYSCCDRPYCNACVGLFANTLVITVTLVTVAGMVGR*

**Fugu Bncra**

…DNLLCYFSPLLEKEVSFKFIATECPPGDLCFKADGRYGNHSALSGRGCMAREACSQTHSIRYKGSVFVMSYSCCDSPYCNSCPGVAAPPFCIAAALLTAALITSPRDVLRGVFSFILE*

<u>Chimeric Bncr sequences</u>

All sequences are derived from zebrafish Bncr except amino acids in bold which are changed to the medaka Bncr sequence and named accordingly.

**Medaka base**

…**ENLHC**LFCPVTSLNSSCAPVVTE**CPPNEMC**YTADGRFGRSSVLFRKG**CMLEKDC**SR SRHQMIRGNNISFSFS**CCDWPYCNRAVALEPFTAMLVAAAVVACSFCLT\***

**Medaka top**

…QGLRC**YYSPVLEKEITFELVVTE**CPVQELC**FKGLGRYGNYTALSARG**CMLRADC**S QVHSLRLLGTVYTMSYS**CCGGHYCNSQPRAEPGGRLLLLLLPAAALTAAGAL*

**Medaka finger 1**

…QGLRC**YYSPVLEKEITFELVVTE**CPVQELCYTADGRFGRSSVLFRKGCMLRADCSRS RHQMIRGNNISFSFSCCGGHYCNSQPRAEPGGRLLLLLLPAAALTAAGAL*

**Medaka finger 2**

…QGLRCLFCPVTSLNSSCAPVVTECPVQELC**FKGLGRYGNYTALSARG**CMLRADCSR SRHQMIRGNNISFSFSCCGGHYCNSQPRAEPGGRLLLLLLPAAALTAAGAL*

**Medaka finger 3**

…QGLRCLFCPVTSLNSSCAPVVTECPVQELCYTADGRFGRSSVLFRKGCMLRADC**SQV HSLRLLGTVYTMSYS**CCGGHYCNSQPRAEPGGRLLLLLLPAAALTAAGAL*

**Medaka fingers 1 + 2**

…QGLRC**YYSPVLEKEITFELVVTE**CPVQELC**FKGLGRYGNYTALSARG**CMLRADCS RSRHQMIRGNNISFSFSCCGGHYCNSQPRAEPGGRLLLLLLPAAALTAAGAL*

**Medaka fingers 1 + 3**

…**ENLHCYYSPVLEKEITFELVVTECPPNEMC**YTADGRFGRSSVLFRKG**CMLEKDCS QVHSLRLLGTVYTMSYSCCDWPYCNRAVALEPFTAMLVAAAVVACSFCLT\***

**Medaka fingers 2 + 3**

…QGLRCLFCPVTSLNSSCAPVVTECPVQELC**FKGLGRYGNYTALSARG**CMLRADC**SQ VHSLRLLGTVYTMSYS**CCGGHYCNSQPRAEPGGRLLLLLLPAAALTAAGAL*

<u>Ancestral state sequences</u>

**Nodes A-D**

…DNLRCYYSPILEKEKTFELIVTECPPDELCFKADGRYGNHSALSARGCMAKKDCGQVHKLRLKGTVYTMSYSCCDWPYCNSQPRAEPGGRLLLLLLPAAALTAAGTL*

**Node E**

…ENLHCYYSPILEKEKTFELIVTECPPNELCFKALGRYGNYTALSARGCMPEKDCSQVHNLRLRGTVYTMSYSCCDWPYCNSQPRAEPGGRLLLLLLPAAALTAAGTL*

**Node F**

…ENLHCYYSPILEKEITFELIVTECPPNELCFKALGRYGNYTALSARGCMLEKDCSQVHSLRLLGTVYTMSYSCCDWPYCNSQPRAEPGGRLLLLLLPAAALTAAGTL*

**Node G**

…DNLRCYYSPILEKEKTFELIVTECPPDELCFKADGRYGNHSALSARGCMAKKDCGQVHKLRFKGTVYTMSYACCDGPYCNSQPRAEPGGRLLLLLLPAAALTAAGTL*

<u>Amino acid substitution sequences</u> Substitution mutations are marked in bold.

**Medaka Bncra I15S**

…ENLHCYYSPVLEKE**S**TFELVVTECPPNEMCFKGLGRYGNYTALSARGCMLEKDCSQV HSLRLLGTVYTMSYSCCDWPYCNRAVALEPFTAMLVAAAVVACSFCLT*

**Medaka Bncra L63R**

…ENLHCYYSPVLEKEITFELVVTECPPNEMCFKGLGRYGNYTALSARGCMLEKDCSQV HSLRL**R**GTVYTMSYSCCDWPYCNRAVALEPFTAMLVAAAVVACSFCLT*

**Medaka Bncra I15S, L63R**

…ENLHCYYSPVLEKE**S**TFELVVTECPPNEMCFKGLGRYGNYTALSARGCMLEKDCSQV HSLRL**R**GTVYTMSYSCCDWPYCNRAVALEPFTAMLVAAAVVACSFCLT*

**Medaka Bncra I15S, A45R**

…ENLHCYYSPVLEKE**S**TFELVVTECPPNEMCFKGLGRYGNYTALS**R**RGCMLEKDCSQV HSLRLLGTVYTMSYSCCDWPYCNRAVALEPFTAMLVAAAVVACSFCLT*

**Medaka Bncra A45R, L63R**

…ENLHCYYSPVLEKEITFELVVTECPPNEMCFKGLGRYGNYTALS**R**RGCMLEKDCSQV HSLRL**R**GTVYTMSYSCCDWPYCNRAVALEPFTAMLVAAAVVACSFCLT*

**Medaka Bncra I15S, A45R, L63R**

…ENLHCYYSPVLEKE**S**TFELVVTECPPNEMCFKGLGRYGNYTALS**R**RGCMLEKDCSQV HSLRL**R**GTVYTMSYSCCDWPYCNRAVALEPFTAMLVAAAVVACSFCLT*

**Zebrafish Bncr S15I**

…QGLRCLFCPVTSLN**I**SCAPVVTECPVQELCYTADGRFGRSSVLFRKGCMLRADCSRSR HQMIRGNNISFSFSCCGGHYCNSQPRAEPGGRLLLLLLPAAALTAAGAL*

**Zebrafish Bncr R63L**

…QGLRCLFCPVTSLNSSCAPVVTECPVQELCYTADGRFGRSSVLFRKGCMLRADCSRSR HQMI**L**GNNISFSFSCCGGHYCNSQPRAEPGGRLLLLLLPAAALTAAGAL*

**Zebrafish Bncr S15I, R63L**

…QGLRCLFCPVTSLN**I**SCAPVVTECPVQELCYTADGRFGRSSVLFRKGCMLRADCSRSR HQMI**L**GNNISFSFSCCGGHYCNSQPRAEPGGRLLLLLLPAAALTAAGAL*

**Zebrafish Bncr S15I, R45A**

…QGLRCLFCPVTSLN**I**SCAPVVTECPVQELCYTADGRFGRSSVLF**A**KGCMLRADCSRSR HQMIRGNNISFSFSCCGGHYCNSQPRAEPGGRLLLLLLPAAALTAAGAL*

**Zebrafish Bncr R45A, R63L**

…QGLRCLFCPVTSLNSSCAPVVTECPVQELCYTADGRFGRSSVLF**A**KGCMLRADCSRSR HQMI**L**GNNISFSFSCCGGHYCNSQPRAEPGGRLLLLLLPAAALTAAGAL*

**Zebrafish Bncr S15I, R45A, R63L**

…QGLRCLFCPVTSLN**I**SCAPVVTECPVQELCYTADGRFGRSSVLF**A**KGCMLRADCSRSR HQMI**L**GNNISFSFSCCGGHYCNSQPRAEPGGRLLLLLLPAAALTAAGAL*

<u>N-glycosylation variant sequences</u>

N-glycosylation site mutations are marked in bold; the three amino acids of one species’ Bncr were changed to the corresponding three amino acids of the other species’ Bncr to either introduce or remove each N-glycosylation consensus sequence.

**Zebrafish Bncr +glyc2**

…QGLRCLFCPVTSLNSSCAPVVTECPVQELCYTADGRFG**NYT**VLFRKGCMLRADCSRS RHQMIRGNNISFSFSCCGGHYCNSQPRAEPGGRLLLLLLPAAALTAAGAL*

**Zebrafish Bncr +glyc2, -glyc3**

…QGLRCLFCPVTSLNSSCAPVVTECPVQELCYTADGRFG**NYT**VLFRKGCMLRADCSRS RHQMIRGN**VYT**FSFSCCGGHYCNSQPRAEPGGRLLLLLLPAAALTAAGAL*

**Zebrafish Bncr +glyc2, -glyc1**

…QGLRCLFCPVTSL**EIT**CAPVVTECPVQELCYTADGRFG**NYT**VLFRKGCMLRADCSRS RHQMIRGNNISFSFSCCGGHYCNSQPRAEPGGRLLLLLLPAAALTAAGAL*

**Zebrafish Bncr +glyc2, -glyc(1+3)**

…QGLRCLFCPVTSL**EIT**CAPVVTECPVQELCYTADGRFG**NYT**VLFRKGCMLRADCSRS RHQMIRGN**VYT**FSFSCCGGHYCNSQPRAEPGGRLLLLLLPAAALTAAGAL*

**Medaka Bncra -glyc2**

…ENLHCYYSPVLEKEITFELVVTECPPNEMCFKGLGRYG**RSS**ALSARGCMLEKDCSQV HSLRLLGTVYTMSYSCCDWPYCNRAVALEPFTAMLVAAAVVACSFCLT*

**Medaka Bncra -glyc2, +glyc1**

…ENLHCYYSPVLEK**NSS**FELVVTECPPNEMCFKGLGRYG**RSS**ALSARGCMLEKDCSQV HSLRLLGTVYTMSYSCCDWPYCNRAVALEPFTAMLVAAAVVACSFCLT*

**Medaka Bncra -glyc2, +glyc3**

…ENLHCYYSPVLEKEITFELVVTECPPNEMCFKGLGRYG**RSS**ALSARGCMLEKDCSQV HSLRLLGT**NIS**MSYSCCDWPYCNRAVALEPFTAMLVAAAVVACSFCLT*

**Medaka Bncra -glyc2, +glyc(1+3)**

…ENLHCYYSPVLEK**NSS**FELVVTECPPNEMCFKGLGRYG**RSS**ALSARGCMLEKDCSQV HSLRLLGT**NIS**MSYSCCDWPYCNRAVALEPFTAMLVAAAVVACSFCLT*

## Data S4. Selection Analyses of Medaka and Zebrafish Bouncer

Selection Analyses of Medaka and Zebrafish Bouncer
| 1 | Positive selection | -1 | Negative selection |  |  |  |  |  |  |  |  |  |
| --- | --- | --- | --- | --- | --- | --- | --- | --- | --- | --- | --- | --- |
| Codon site in alignment | MEME | MEME number of branches under selection | FUBAR | FEL all | FEL Danio branch | FEL Oryzias branch | FEL Contrast Danio vs. Oryzias | BUSTED all | Danio rerio AA in alignment | Oryzias latipes AA in alignment | Danio rerio site | Oryzias latipes site |
| 1 | 0 | 0 | -1 | -1 | 0 | 0 | 0 | 0 | Q | E | 1 | 1 |
| 2 | 0 | 0 | -1 | -1 | 0 | 0 | 0 | 0 | G | N | 2 | 2 |
| 3 | 1 | 2 | 0 | 0 | 0 | 0 | 0 | 0 | L | L | 3 | 3 |
| 4 | 1 | 1 | -1 | -1 | 0 | 0 | 0 | 0 | R | H | 4 | 4 |
| 5 | 0 | 1 | -1 | -1 | -1 | 0 | 0 | 0 | C | C | 5 | 5 |
| 6 | 0 | 0 | 0 | 0 | 0 | 0 | 0 | 0 | L | Y | 6 | 6 |
| 7 | 0 | 1 | 0 | 0 | 0 | 0 | 0 | 0 | F | Y | 7 | 7 |
| 8 | 0 | 0 | -1 | -1 | -1 | 0 | 0 | 0 | C | S | 8 | 8 |
| 9 | 0 | 1 | -1 | -1 | -1 | 0 | 0 | 0 | P | P | 9 | 9 |
| 10 | 0 | 0 | 0 | 0 | 0 | 0 | 0 | 0 | V | V | 10 | 10 |
| 11 | 0 | 0 | 0 | 0 | 0 | 0 | 0 | 0 | T | L | 11 | 11 |
| 12 | 0 | 0 | 0 | 0 | 0 | 0 | 0 | 0 | S | E | 12 | 12 |
| 13 | 0 | 0 | 0 | 0 | 0 | 0 | 0 | 0 | - | - | 12 | 12 |
| 14 | 0 | 1 | 0 | 0 | 0 | 0 | 0 | 0 | - | - | 12 | 12 |
| 15 | 0 | 0 | -1 | -1 | 0 | 0 | 0 | 0 | L | K | 13 | 13 |
| 16 | 0 | 0 | 0 | 0 | 0 | 0 | 0 | 0 | N | E | 14 | 14 |
| 17 | 1 | 6 | 0 | 1 | 1 | 1 | 0 | 0 | S | I | 15 | 15 |
| 18 | 0 | 1 | -1 | -1 | 0 | 0 | 0 | 0 | S | T | 16 | 16 |
| 19 | 0 | 3 | -1 | -1 | -1 | 0 | 0 | 0 | C | F | 17 | 17 |
| 20 | 0 | 0 | 0 | 0 | 0 | 0 | 0 | 0 | A | E | 18 | 18 |
| 21 | 0 | 0 | -1 | -1 | 0 | 0 | 0 | 0 | P | L | 19 | 19 |
| 22 | 0 | 4 | 0 | -1 | -1 | 0 | 0 | 0 | V | V | 20 | 20 |
| 23 | 0 | 0 | 0 | 0 | 0 | 0 | 0 | 0 | V | V | 21 | 21 |
| 24 | 0 | 0 | -1 | -1 | 0 | 0 | 0 | 0 | T | T | 22 | 22 |
| 25 | 0 | 2 | -1 | -1 | -1 | 0 | 0 | 0 | E | E | 23 | 23 |
| 26 | 0 | 0 | -1 | -1 | -1 | 0 | 0 | 0 | C | C | 24 | 24 |
| 27 | 0 | 0 | -1 | -1 | -1 | 0 | 0 | 0 | P | P | 25 | 25 |
| 28 | 0 | 0 | -1 | -1 | 0 | 0 | 0 | 0 | V | P | 26 | 26 |
| 29 | 0 | 0 | -1 | 0 | -1 | 0 | 0 | 0 | Q | N | 27 | 27 |
| 30 | 0 | 0 | -1 | 0 | 0 | 0 | 0 | 0 | E | E | 28 | 28 |
| 31 | 0 | 0 | 0 | 0 | 0 | 0 | 0 | 0 | L | M | 29 | 29 |
| 32 | 0 | 0 | 0 | 0 | 0 | 0 | 0 | 0 | - | - | 29 | 29 |
| 33 | 0 | 0 | -1 | -1 | 0 | 0 | 0 | 0 | C | C | 30 | 30 |
| 34 | 0 | 0 | -1 | -1 | -1 | -1 | 0 | 0 | Y | F | 31 | 31 |
| 35 | 0 | 0 | -1 | -1 | -1 | 0 | 0 | 0 | T | K | 32 | 32 |
| 36 | 0 | 0 | 0 | 0 | 0 | 0 | 0 | 0 | - | - | 32 | 32 |
| 37 | 0 | 0 | 0 | 0 | 0 | 0 | 0 | 0 | - | - | 32 | 32 |
| 38 | 0 | 0 | 0 | 0 | 0 | 0 | 0 | 0 | - | - | 32 | 32 |
| 39 | 0 | 0 | 0 | 0 | 0 | 0 | 0 | 0 | - | - | 32 | 32 |
| 40 | 0 | 5 | -1 | -1 | -1 | 0 | 0 | 0 | A | G | 33 | 33 |
| 41 | 1 | 1 | -1 | -1 | 0 | 0 | 0 | 0 | D | L | 34 | 34 |
| 42 | 0 | 3 | 0 | 0 | 0 | 0 | 0 | 0 | G | G | 35 | 35 |
| 43 | 0 | 2 | 0 | 0 | 0 | 0 | 0 | 0 | R | R | 36 | 36 |
| 44 | 0 | 0 | -1 | -1 | 0 | 0 | 0 | 0 | F | Y | 37 | 37 |
| 45 | 0 | 0 | -1 | -1 | -1 | 0 | 0 | 0 | G | G | 38 | 38 |
| 46 | 0 | 0 | -1 | -1 | 0 | 0 | 0 | 0 | R | N | 39 | 39 |
| 47 | 1 | 2 | 0 | 0 | 0 | 0 | 0 | 0 | S | Y | 40 | 40 |
| 48 | 0 | 0 | -1 | -1 | 0 | 0 | 0 | 0 | S | T | 41 | 41 |
| 49 | 0 | 0 | 0 | -1 | 0 | 0 | 0 | 0 | V | A | 42 | 42 |
| 50 | 0 | 0 | -1 | -1 | -1 | 0 | 0 | 0 | L | L | 43 | 43 |
| 51 | 0 | 0 | -1 | -1 | 0 | 0 | 0 | 0 | F | S | 44 | 44 |
| 52 | 0 | 0 | -1 | 0 | 0 | 0 | 0 | 0 | R | A | 45 | 45 |
| 53 | 0 | 0 | -1 | -1 | 0 | 0 | 0 | 0 | K | R | 46 | 46 |
| 54 | 0 | 0 | -1 | -1 | -1 | 0 | 0 | 0 | G | G | 47 | 47 |
| 55 | 0 | 0 | -1 | -1 | -1 | 0 | 0 | 0 | C | C | 48 | 48 |
| 56 | 0 | 1 | 0 | 0 | 0 | 0 | 0 | 0 | M | M | 49 | 49 |
| 57 | 0 | 0 | -1 | 0 | 0 | 1 | 0 | 0 | L | L | 50 | 50 |
| 58 | 0 | 0 | 0 | 0 | 0 | 0 | 0 | 0 | R | E | 51 | 51 |
| 59 | 0 | 0 | -1 | -1 | 0 | 0 | 0 | 0 | A | K | 52 | 52 |
| 60 | 1 | 6 | 0 | 0 | 0 | 0 | 0 | 0 | D | D | 53 | 53 |
| 61 | 0 | 0 | -1 | -1 | -1 | 0 | 0 | 0 | C | C | 54 | 54 |
| 62 | 0 | 0 | 0 | 0 | 0 | 0 | 0 | 0 | S | S | 55 | 55 |
| 63 | 0 | 0 | 0 | 0 | 0 | 0 | 0 | 0 | R | Q | 56 | 56 |
| 64 | 0 | 0 | -1 | 0 | 0 | 0 | 0 | 0 | S | V | 57 | 57 |
| 65 | 0 | 0 | 0 | 0 | 0 | 0 | 0 | 0 | R | H | 58 | 58 |
| 66 | 0 | 0 | -1 | -1 | 0 | 0 | 0 | 0 | H | S | 59 | 59 |
| 67 | 0 | 0 | -1 | -1 | 0 | 0 | 0 | 0 | Q | L | 60 | 60 |
| 68 | 0 | 0 | -1 | 0 | 0 | 0 | 0 | 0 | M | R | 61 | 61 |
| 69 | 0 | 0 | 0 | 0 | 0 | 0 | 0 | 0 | I | L | 62 | 62 |
| 70 | 0 | 1 | -1 | -1 | 0 | 1 | Oryzias<br>positive | 1 | R | L | 63 | 63 |
| 71 | 0 | 0 | -1 | -1 | -1 | 0 | 0 | 0 | G | G | 64 | 64 |
| 72 | 0 | 0 | -1 | -1 | 0 | 0 | 0 | 0 | N | T | 65 | 65 |
| 73 | 0 | 0 | -1 | -1 | -1 | 0 | 0 | 0 | N | V | 66 | 66 |
| 74 | 0 | 0 | 0 | 0 | 0 | 0 | 0 | 0 | I | Y | 67 | 67 |
| 75 | 0 | 0 | 0 | 0 | 0 | 0 | 0 | 0 | S | T | 68 | 68 |
| 76 | 0 | 0 | 0 | 0 | 0 | 0 | 0 | 0 | F | M | 69 | 69 |
| 77 | 0 | 0 | 0 | -1 | 0 | 0 | 0 | 0 | S | S | 70 | 70 |
| 78 | 0 | 0 | 0 | 0 | 0 | 0 | 0 | 0 | F | Y | 71 | 71 |
| 79 | 0 | 0 | 0 | 0 | 0 | 0 | 0 | 0 | S | S | 72 | 72 |
| 80 | 0 | 0 | -1 | -1 | 0 | 0 | 0 | 0 | C | C | 73 | 73 |
| 81 | 0 | 0 | -1 | -1 | -1 | 0 | 0 | 0 | C | C | 74 | 74 |
| 82 | 0 | 0 | -1 | -1 | 0 | 0 | 0 | 0 | G | D | 75 | 75 |
| 83 | 0 | 2 | 0 | 0 | 0 | 0 | 0 | 0 | G | W | 76 | 76 |
| 84 | 0 | 0 | -1 | -1 | 0 | 0 | 0 | 0 | H | P | 77 | 77 |
| 85 | 0 | 0 | 0 | 0 | 0 | 0 | 0 | 0 | Y | Y | 78 | 78 |
| 86 | 0 | 0 | -1 | -1 | -1 | 0 | 0 | 0 | C | C | 79 | 79 |
| 87 | 0 | 0 | -1 | -1 | -1 | 0 | 0 | 0 | N | N | 80 | 80 |

